# Fixation and effective size in a haploid-diploid population with asexual reproduction

**DOI:** 10.1101/2020.09.17.295618

**Authors:** Kazuhiro Bessho, Sarah P. Otto

**Affiliations:** Saitama Medical University, 38 Morohongo Moroyama-machi, Iruma-gun, Saitama 350-0495, Japan; Department of Zoology, The University of British Columbia, Vancouver, BC, V6T 1Z4, Canada

**Keywords:** haploid-diploid life cycle, Wright-Fisher model, fixation probability, variance effective population size, class reproductive value

## Abstract

The majority of population genetic theory assumes fully haploid or diploid organisms with obligate sexuality, despite complex life cycles with alternating generations being commonly observed. To reveal how natural selection and genetic drift shape the evolution of haploid-diploid populations, we analyze a stochastic genetic model for populations that consist of a mixture of haploid and diploid individuals, allowing for asexual reproduction and niche separation between haploid and diploid stages. Applying a diffusion approximation, we derive the fixation probability and describe its dependence on the reproductive values of haploid and diploid stages, which depend strongly on the extent of asexual reproduction in each phase and on the ecological differences between them.

**Highlight:**

- Classical models consider fully haploid or diploid populations
- We model haploid-diploid life cycles allowing for asexual reproduction
- We obtain the fixation probability of alleles subject to selection and drift
- Reproductive values of haploid and diploid stages shape their evolution

## 1. Introduction

Sexual reproduction in eukaryotes generally consists of an alternation of generations, where meiosis halves the number of chromosomes to produce haploids and syngamy brings together haploid gametes to produce diploids. The extent of development in each ploidy phase varies substantially (Bell 1982; 1994). In diplontic organisms, at one extreme, development and growth occur only in the diploid phase, as is observed in most animals. Haplontic organisms, at the other extreme, undergo mitotic growth only in the haploid stage, as is seen in some green algae. In between these extremes, many terrestrial plants, macroalgae, and fungi exhibit both haploid and diploid growth (haploid-diploid life cycles). These stages are typically free living in macroalgae, with either macroscopically similar (isomorphic) or distinct (heteromorphic) forms in the haploid and diploid stage (Raper and Flexer 1970; Wilson 1981; Mable and Otto 1998; Coelho 2007).

To explain variation in life cycles, several theoretical models have analyzed the deterministic dynamics of a modifier allele that alters the time spent in haploid and diploid phases (e.g., Perrot et al. 1991; Otto and Goldstein 1992; Goldstein 1992; Otto 1994; Orr and Otto 1994; Jenkins and Kirkpatrick 1995; Otto and Marks 1996; Scott and Rescan 2017).However, there are some gaps between these models and the complexities seen in many haploid-diploid species. For example, these models often treat haploid and diploid individuals as ecologically equivalent, despite the frequent observation of niche differences and seasonal shifts in prevalence (e.g., Slocum 1980; Dethier 1981). Most of these models also assume obligate sexuality (but see Otto and Marks 1996), despite asexuality being frequently observed among haploid-diploid species (“asexual looping”). Furthermore, while several models have explored how haploid-diploid life cycles might evolve, the impact of haploid-diploid life cycles on evolutionary processes remains underexplored (see, e.g., Bessho and Otto 2017 on the impact on fixation probabilities and Immler et al. 2012 on the maintenance of variation).

Here we contribute to evolutionary theory for haploid-diploid populations by calculating the fixation probability of mutations using a stochastic genetic model. This builds upon our previous work (Bessho and Otto 2017) by accounting for asexual looping and niche differences between ploidy phases, both of which are common in macroalgae (Bell 1982; de Wreede and Klinger 1988; Hawkes 1990). Haploid and diploid phases often differ physiologically, and even isomorphic haploids and diploids may differ ecologically (Hannach and Santelices 1985; Destombe et al. 1993; Dyck and de Wreede 2006; Thornber et al., 2006; Vieira et al. 2018). We therefore explore different forms of density dependence, acting either globally on the total population size (as in Bessho and Otto 2017) or locally on the population size of haploids and diploids separately (Figure 1). We show that the fate of a mutation depends strongly on the reproductive values of haploids and diploids, which in turn depend on the extent of asexual reproduction and ecological differences between the phases.

**Fig. 1.**
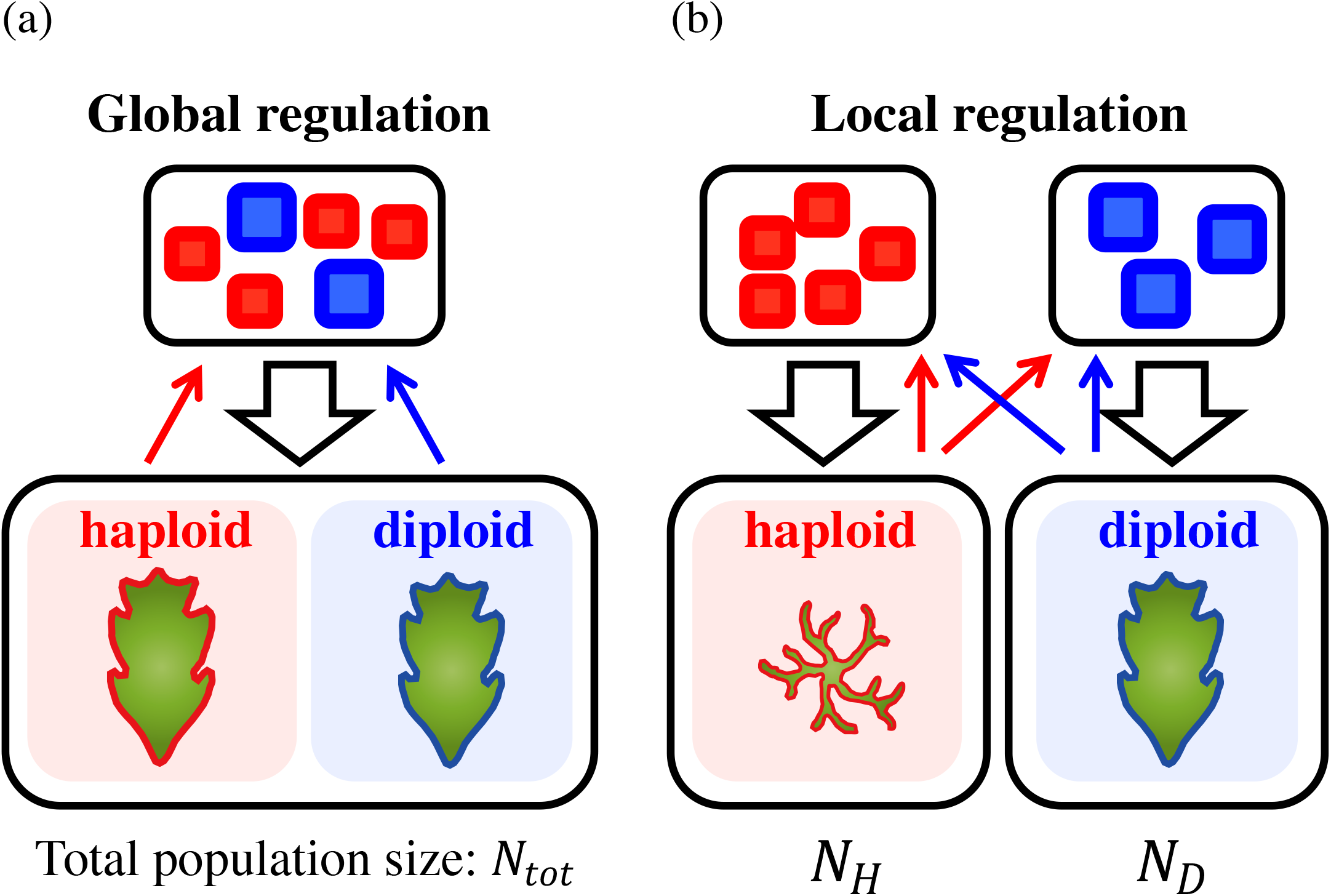
An illustration of the haploid-diploid models. (a) In the global regulation model, both haploids and diploids occupy the same habitat and density dependence holds the total population size *N*_*tot*_ constant. (b) In the local regulation model, each ploidy stage occupies a different habit, therefore density dependence regulates the population size of haploids (*N*_*H*_) and diploids (*N*_*D*_) separately.

## 2. Model

In Bessho and Otto (2017), we calculated the fixation probabilities by tracking the dynamics of a resident allele (*R*) and a mutant allele (*M*) in haploid and diploid individuals, using both a Wright-Fisher and a Moran model. In that model, reproduction was obligately sexual, individuals were ecologically equivalent, and the total population size was held constant (global density dependence). Below, we calculate the fixation probability by first considering asexual reproduction in each phase, assuming that haploids and diploids are ecologically equivalent (global population regulation), and then determine how these results are affected by niche differences (local population regulation that is ploidy specific).

### 2.1 Haploid-diploid Wright-Fisher model with global regulation and asexual looping

Let *x*_(*GT*)_ (*t*) be a random variable that represents the number of individuals of a particular genotype (“*GT”*) at time *t*, involving the resident (*R*) and mutant alleles (*M*) (e.g., *GT* = *RM* for a heterozygous diploid and *GT* = *M* for a mutant haploid). Let *x*_(GT)_ (*t*) represent a particular outcome of this random variable. In the global regulation model, we assume a constant population, *x*_*R*_+ *x*_*M*_ + *x*+ *x*_*RR*_ + *x*_*RM*_ + *x*_*M*_ + *x*_*MM*_ =*N*_*tot*_, that is strictly regulated regardless of the ploidy of the individuals.

The reproductive output and the degree of asexuality are characterized by *w*_(*GT*)_ and *a*_*H*_ for haploids [(*GT*) = *R* or *M*] and *w*_(*GT*)_ and *a*_*D*_ for diploids [(*GT*) = *RR, RM*, and *MM*]. Specifically, diploid individuals produce (1 − *a*_*D*_ *w*_(*GT*)_ haploid spores (sexual reproduction) and *a*_*D*_ *w*_(*GT*)_ diploid offspring (asexual loop). Similarly, haploids produce (1 − *a*_*H*_) *w*_(*GT*)/2_ female gametes (sexual reproduction) and *a*_*H*_ *w*_(*GT*)_ haploid offspring (asexual loop), where we assume that the species is monoecious and invests equal resources in male and female gametes. During syngamy, we assume that male gametes are not limiting, that mating is random, and that female gametes are successfully fertilized with male gametes, at a rate *f*_(*GT*)_ [(*GT*) = *R* or *M*], becoming diploid zygotes. For clarity, we describe the model with non-overlapping generations, although we note that overlapping generations can be considered by including surviving adults in the counts of asexual offspring (*a*_*D*_ *w*_(*GT*)_ *a*_*H*_*w*_(*GT*))_

We define the selection coefficient (*S*_(GT)_) and the degree of dominance (*h*) acting upon the mutant allele such that: 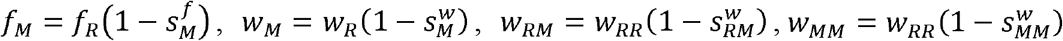, and 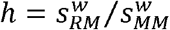. To perform the diffusion approximation, we assume that selection is weak, 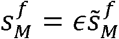 and 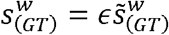, where ϵ is a small parameter (parameter ϵ is order *O* (1/*N*_*tot*_)).

### 2.2 Haploid-diploid Wright-Fisher model with local regulation and asexual looping

We then consider the case where density dependence regulates haploid and diploid populations separately, which may occur if they have different resource needs or utilize different habitats or microhabitats (for short-hand, we refer to this case as “local regulation”). More specifically, we assume that the population size of haploids and diploids is separately regulated and remains constant *N*_*H*_ and *N*_*D*_ (*x*_*R*_ − *xM*= *N*_*H*_ and *x*_*RR*_ − *x*_*RM*_ − *x*_*MM*_=*N*_*D*_), respectively. We set 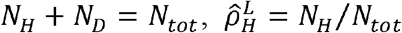 and 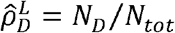, which will then allow us to and remains compare the results of local and global regulation. Holding population sizes constant is assumed strictly for mathematical convenience but may be reasonable for populations whose sizes are strongly regulated by the availability of appropriate habitat.

## 3. Fixation probability in a haploid-diploid population

### 3.1 Fixation probability in the global regulation model

The fixation probability in a haploid-diploid population can be derived using a diffusion approximation (Bessho and Otto 2017), but doing so requires that we approximate the dynamics to reduce the dimensionality from four variables (*x*_*R*_, *x*_*m*_, *x*_*RR*_, *x*_*RM*_, *x*_*MM*_, which sum to *N*_*tot*_) down to one. We do so by using a separation of time scales, deriving the first and second moments of the mutant allele frequency. Specifically, we transform the number of individuals of each genotype,*S*_(*GT*)_, into new variables that allow us to separate the slower evolutionary dynamics and the faster ecological dynamics (Appendix A):

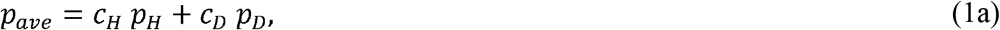

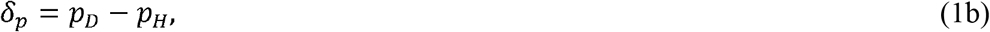

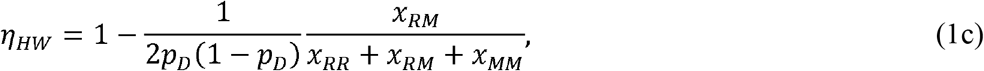

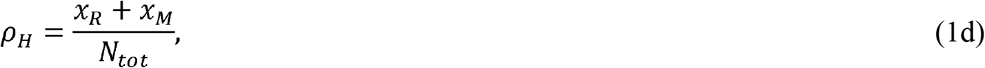

Where *p*_*ave*_ indicates the average allele frequency of haploids and diploids weighted by the class reproductive values (*C*_*H*_ and *C*_*D*_, where *C*_*H*_ − *CD* = 1, see next paragraph), *δ*_*p*_ indicates the where indicates the average allele frequency of haploids and diploids weighted by the class difference in allele frequencies between haploids and diploids, *η*_*HW*_ indicates the departure from the Hardy-Weinberg equilibrium in diploids, and *ρ*_H_ indicates the frequency of haploids in the population. Within these equations, the frequencies of mutant alleles in haploids and diploids are *P*_*H*_ = *x*_*M*_/ (*x*_*R*_ + *x*_*M*_) and 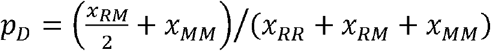 As similar variables are used in the model with local population regulation, we use superscripts to indicate the form of population regulation (“*Model*” is *G* for global or *L* for local regulation).

The class reproductive values of haploids and diploids are defined as follows. In linear models, “reproductive value” is a measure of the expected fraction of the population in the long-term future that descends from an individual of a particular type (e.g., age or stage class). Class reproductive values, as defined by Taylor (1990) and Rousset (2004, p.153), scale these individual reproductive values up to the whole population of each class (i.e., the product of the individual reproductive values times the class size). In the models considered here, the dynamics are non-linear because of competition for resources (*N*_*tot*_.) Nevertheless, we can approximate reproductive values by assuming that the population is near equilibrium with only resident alleles and by holding the strength of competition constant (Appendix A, Supplementary *Mathematica* file; Supplementary parts). Doing so, we find that the class reproductive values of haploids and diploids, expressed as proportions that sum to one, are:

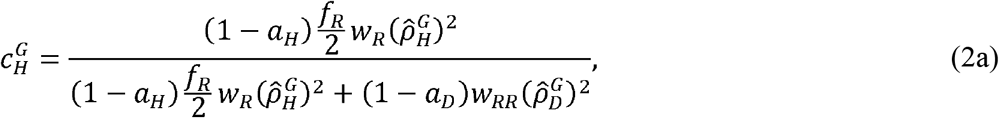

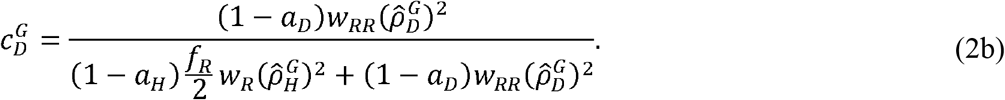

where 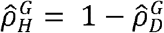 is the equilibrium frequency of haploids in the global model (Eq. A.4). As a special case of interest, when populations are purely sexual (*a*_*H*_ =*a*_*D*_ = 0), we can plug the equilibrium for 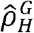 from Eq. (A.4) into (2) and show that 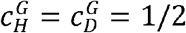 As discussed in section 3.4 below, increasing the degree of asexual reproduction of haploids tends to increase their reproductive value, but there are exceptions when diploids are more fertile and predominantly asexual. Similarly, increasing the extent of asexual reproduction among diploids tends to increase the reproductive value of diploids, except when haploids are more fertile and predominantly asexual. These exceptions occur because passage through the more productive ploidy level (via sexual reproduction) then becomes essential for individuals of the less productive ploidy to contribute substantially to future generations.

As discussed in Appendix A (see also Bessho and Otto 2017), equations (2) provide the only weights that allow ecological and evolutionary time scales to be separated when calculating the average allele frequency in equation (1a), which is why we take that to be the evolutionarily relevant average. Although one might initially think that diploids should count twice as much because they contain two allele copies and that the evolutionarily relevant average allele frequency would depend on the population sizes of haploids and diploids, having a strict alternation of generations in a fully sexual population (*a*_*H*_ =*a*_*D*_ = 0) ensures that haploids and diploids contribute equally to long-term future generations, so that their reproductive values are equal and the evolutionarily relevant average allele frequency is *pave* = (1/2)*p*_*H*_ + (1/2) *p*_*D*_ (Bessho and Otto 2017).

As with our previous model, we can track the slow evolutionary dynamics for the expected change in average allele frequency *p*_*ave*_ under weak selection, once the fast ecological dynamics have stabilized, as which point we can show that there are similar allele frequencies in haploids and diploids (*δ*_*p*_ ≈ = 0) diploids are approximately at Hardy-Weinberg equilibrium (*η*_*HW*_ ≈ = 0), and the ratio of haploids is similar among mutant and resident genotypes 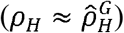 (Appendix A, Supplementary *Mathematica* file; Diffusion approximation for global (local) regulation model). Furthermore, to leading order, the second moment of change in allele frequency is equal to the neutral case and can be derived in the diffusion limit (*N*_*tot*_ → ∞).

Given a single variable, *p*_*ave*_, changing slowly over evolutionary time, we can then use standard diffusion methods to calculate the fixation probability of a mutation in a haploid-diploid population, 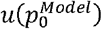, where “*Model*” is *G* for global and *L* for local (considered in the next section). The diffusion is a function of the first and second moments of change in the mutant allele frequency, *m* ^*Model*^ (*p*_*ave*_) and *v* ^*Model*^ (*p*_*ave*_), both measured in time units of *N*_*tot*_ in the generations:

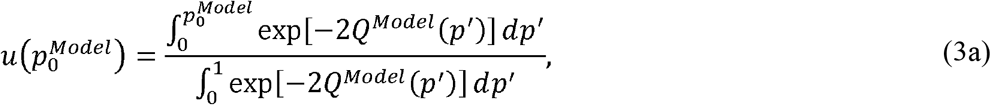

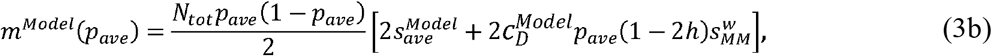

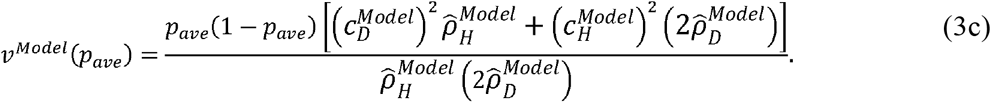

where 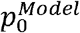 is the initial allele frequency of mutants (we focus on the case with a single initial mutant allele, 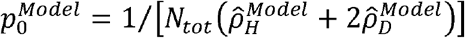 and *Q* ^*Model*^ (*p*) = *∫* (*m* ^*Model*^ (*p*) / *v* ^*Model*^ (*p*))*dp* We note that Eq (3a) is obtained from the backward equation for the fixation probability (Bessho and Otto 2017) and Eqs. (3b) and (3c) are derived in Appendix A. For the global regulation model, the average selection acting upon rare mutant alleles across haploid and diploid stages, 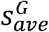, can be calculated from the first moment equation (see Supplementary *Mathematica* file; Diffusion approximation for the global (local) regulation model) and equals:

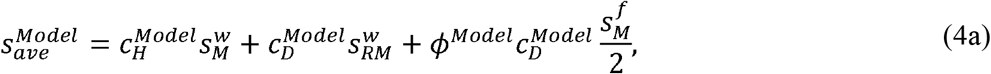

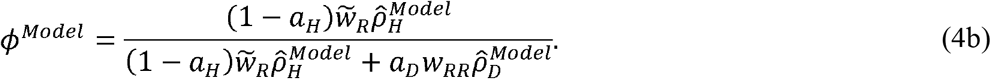

where 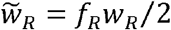 is the fertility of haploids considering the cost of sex. As we will see later, *Ø*^*Model*^ this equation is valid for local regulation model. The term *ØModel* indicates the fraction of the diploids in the next generation that are sexually produced (from the union of haploid-produced gametes) rather than asexually produced by diploid parents. With obligately sexual haploid-diploids(*a*_*H*_ = *a*_*D*_ = 0), where 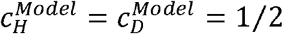 and *Ø*^*Model*^ = 1), these results coincide with those of Bessho and Otto (2017).

### 3.2 Fixation probability in the local regulation model

We next derive the fixation probability in a haploid-diploid population when density dependence regulates haploid and diploid populations separately (Figure 1b), by again applying a transformation of variables and separation of time scales. For the local regulation model, the appropriate weights for the average allele frequency are similar to the global regulation model, where now the class reproductive values, expressed as proportions, are:

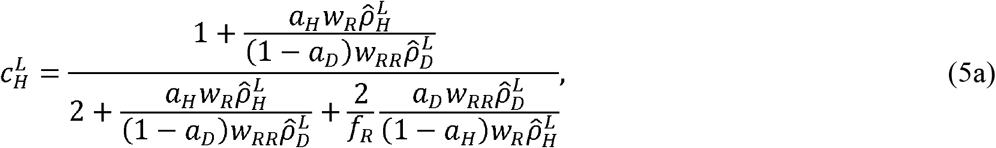

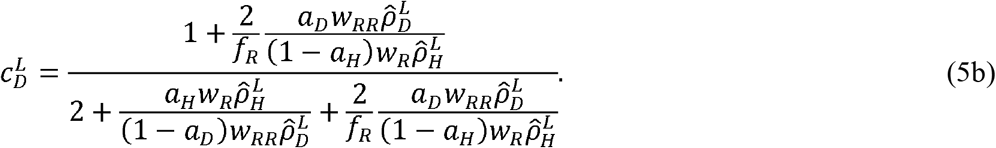

After applying a separation of time scales and conducting a diffusion approximation, we conclude that the solution for the fixation probability in a haploid-diploid population, Eqs. (3), remains valid for the local regulation model (Supplementary *Mathematica* file; Diffusion approximation for the local regulation model), with the average selection coefficient now being given by Eqs. (5).

### 3.3 Effective genetic parameters

Using the first and second moments of change in allele frequency, we derive effective genetic parameters to compare our results to the dynamics found in the classical model for fully haploid or fully diploid organisms (Bessho and Otto 2017). More specifically, we define the effective selection coefficient (*S*_e_), dominance coefficient (*h*_e_), and effective population size (*N*_e_) that would result in the same expected change in allele frequency and variance as in the classical diploid model of selection.

For selection, the diploid model is: Δ *p*_*ave*_ *= S*_e_ *p*_*ave*_ (1− *p*_*ave*_)[*h*_e_ + 1 − 2*he*) *p*_*ave*_] (Crow and Kimura 1970; Bessho and Otto 2017). Because this equation depends on the allele frequency in the same way as Eq. (3b), we can find the effective and dominance selection coefficient from Δ*p*_*ave*_ /[*p*_*ave*_ (1− *p*_*ave*_)]=*S*_*e*_ *h*_*e*_ when *p*_*ave*_ =0 and Δ*p*_*ave*_ /[*p*_*ave*_ (1− *p*_*ave*_)]=*S*_*e*_ (1 − *h*_*e*_) when *p*_*ave*_*=*1, yielding:

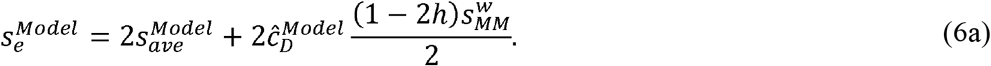

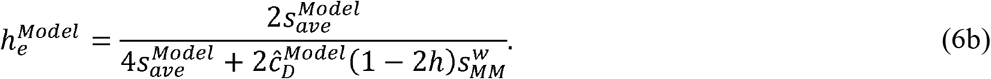

When the mutation is additive(*h*=1/2), these effective parameters are 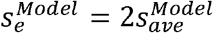 and 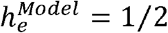.

We next derive the variance effective population size (Crow and Kimura 1970) by equating the one generation change in variance (Eq. 3c divided by the time scale,*N*_*tot*_) to the variance in allele frequency expected in the classical Wright-Fisher model, *p*_*ave*_ (1 − *p*_*ave*_)/2*N*_*e*_with *N*_*e*_ diploid individuals, obtaining:

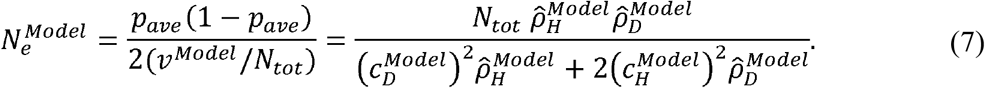

Plugging these effective parameters into the formula from the fixation probability in the classical diploid Wright-Fisher model (Kimura 1957; 1962; Crow and Kimura 1970, p. 427), the fixation probability in a haploid-diploid population given by Eq. 3a can be expressed as:

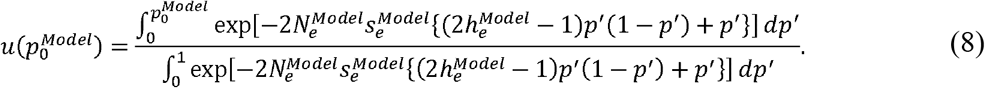

Assuming an initially rare and additive mutation(*h* =1/2) with weak positive selection in a large population (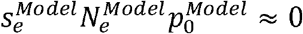 and 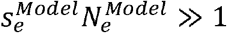), we obtain the classic approximation, 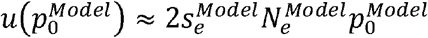, which upon substituting from Eq. 7 yields:

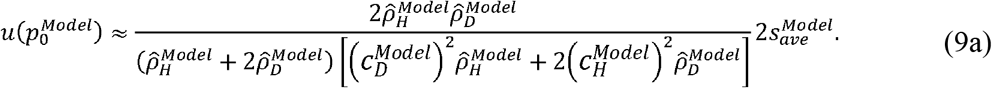

For example, because haploids and diploids have the same reproductive values in the obligately sexual case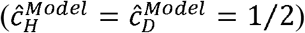, we obtain:

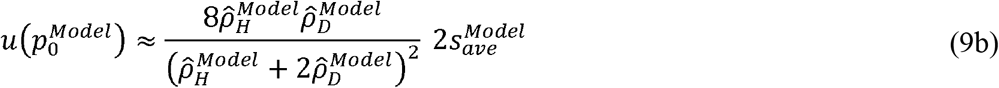

(Eq. 13a in Bessho and Otto 2017), or simply 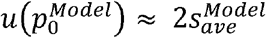 if haploid and diploid population sizes are equal in terms of number of chromosomes 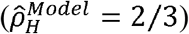.

In the next three sections, we explore the implications of these results for the evolution of haploid-diploid populations.

### 3.4 Effective selection in a haploid-diploid population

The strength of selection averaged across haploids and diploids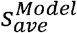, plays a key role in the evolution of haploid-diploid populations. When a mutation is rare, both the rate of change in allele frequency (Eq. 3b) and the approximate fixation probability (Eq. 10a) are proportional to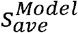. We thus begin by exploring how 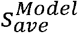 varies as we alter the amount of asexual reproduction in haploid and diploid phases. We focus on the case where the mutation does not affect fertilization success 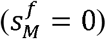, so that the average selection becomes.

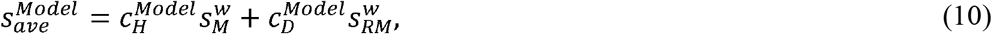

in both global and local regulation models (Eqs. (4) and (6)).

The relative evolutionary importance of selection in the haploid and diploid phases is thus determined by the class reproductive values, 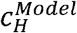 and 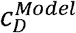 (where 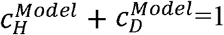). Figures 2 (global regulation) and 3 (local regulation) illustrate the proportional reproductive value of haploids, 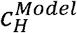, as a function of the degree of asexual reproduction in haploids (x-axis) and diploids (ranging from 0.05 in red to 0.95 in blue). With global regulation, the frequency of haploidy within the population, 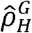 (given by Eq. A.4), varies with the parameters (see inset graphs in Figure 2), rising with the frequency of haploid asexuality (x-axis in inset) but declining with more asexuality in diploids (from red to blue).

**Fig. 2.**
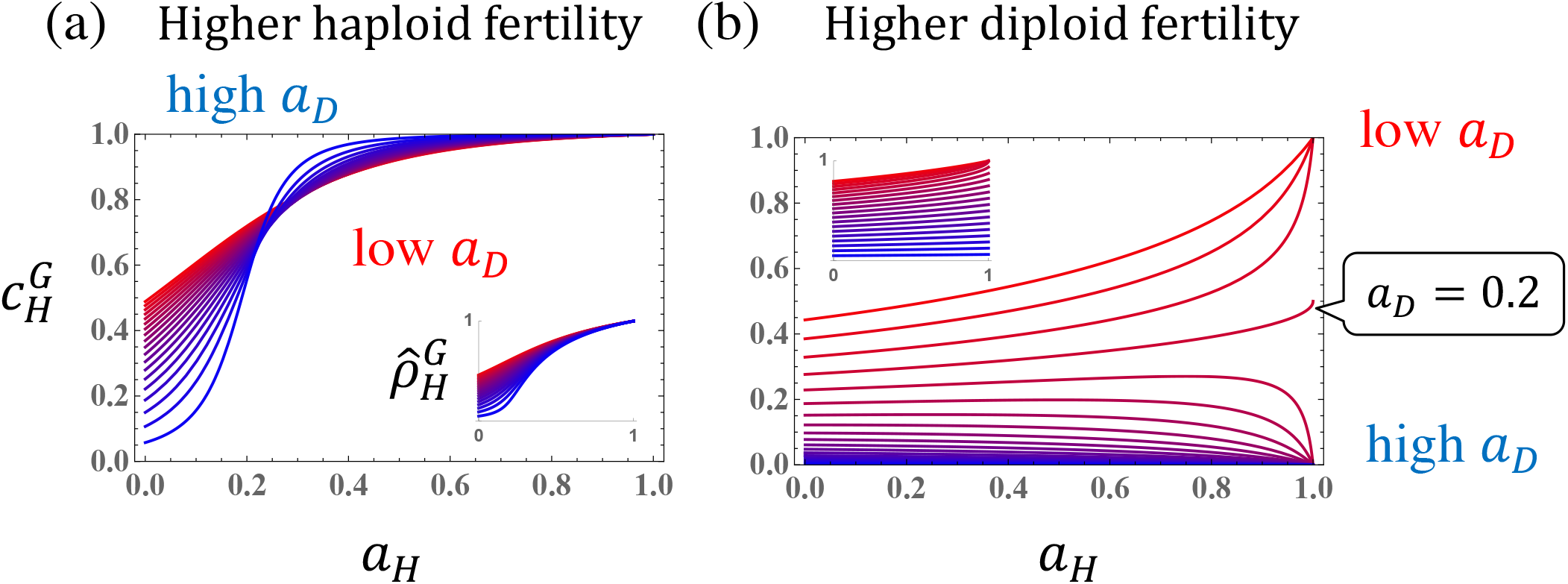
Class reproductive value of haploids in the global regulation model,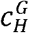. Curves show 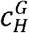 as a function of the degree of haploid asexuality, *a*_*H*_ (x-axis), with the degree of diploid asexuality ranging in color from *a*_*D*_ = 0.05 (red) to 0.95 (blue) in increments of 0.05. Other parameters are set as: (a) *f*_*R*_ = 0.5, *w*_*R*_ = 5000, *w*_*RR*_ *=* 1000, (b) *f*_*R*_ = 0.5, *w*_*RR*_ = *1000 w*_*RR*_ *=* 5000. The resulting frequency of haploids, 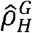 (Eq. A.4), is shown in the inset plots. Note that if haploids are strictly asexual (*a*_*H*_ =1), diploids persist only when *a*_*D*_*W*_*RR*_ > *W*_*R*_ (i.e., when *a*_*D*_ > 0.2 in panel b), which accounts for the switch in behavior observed in this panel.

By contrast, with local regulation, the frequency of haploidy is held fixed by the strict density dependent competition that we have assumed (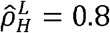 in Figure 3(a)(b) and 0.3 in 3(c)(d)). In the left panels, haploids have a higher fertility (*w*_*R*_/*w*_*RR*_ = 5), leading to a higher haploid reproductive value, 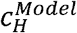, especially with local regulation when haploids are also more common (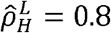 in Figure 3a). In the right panels, diploids have a higher fertility (*w*_*RR*_ /*w*_*R*_ = 5)), leading to a lower haploid reproductive value, especially when haploids are rare (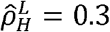 in Figure 3d).

**Fig. 3.**
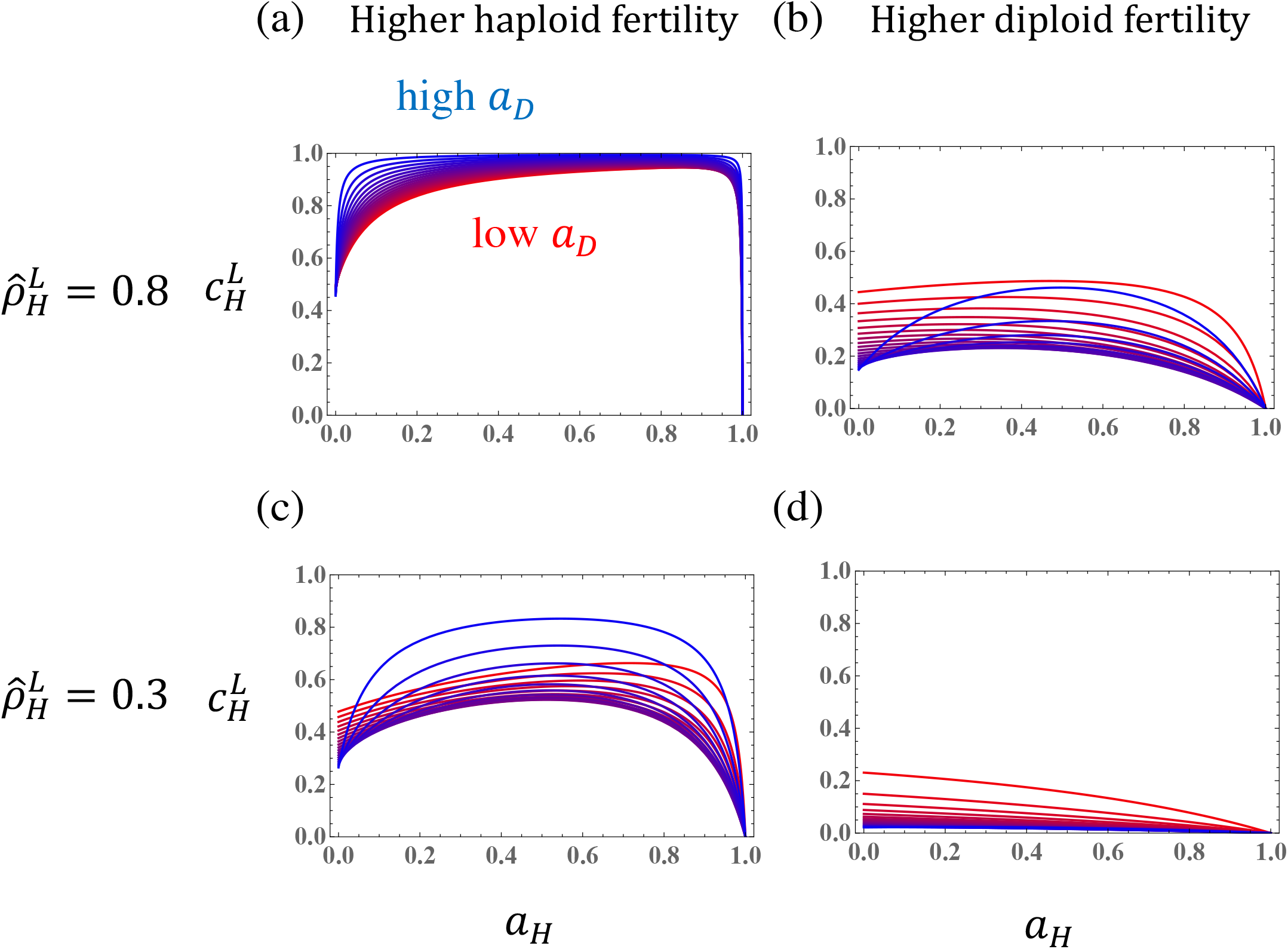
Proportional reproductive value of haploids in the local regulation model, 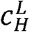. Parameters are the same as Fig. 2, except that haploids are held fixed at a frequency of (a)(b) 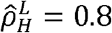 or (c)(d) 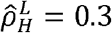. We consider the case when (a)(c) haploid fertility is larger than diploid *w*_*R*_ *=* 5000, *w*_*RR*_ *=* 1000 and when (b)(d) diploid fertility is larger than haploid *w*_*R*_ *=* 1000, *w*_*RR*_ *=* 5000.

When haploids are primarily sexual (*a*_*H*_ ≈ 0), increasing asexuality of the haploid stage typically causes the reproductive value of haploids to rise, unless diploids are fitter and more frequent (Figure 3d and blue curves in Figure 2b). At the other extreme, the reproductive value of haploids typically plummets to zero as haploid reproduction becomes primarily asexual (*a*_*H*_ ≈1) while diploids remain sexual, particularly with local regulation (Figure 3), because haploids then act as a genetic “sink” contributing little to the diploid sub-population. This downward trend when haploids are predominantly asexual is also seen with global regulation if diploids are more fit (Figure 2b), except when the diploid population does not sustain itself and goes extinct, which occurs when *a*_*D*_ < 0.2 and *a*_*H*_ =1. The net result can thus be non-monotonic (purple curves with0.2 < *a*_*D*_ < in Figure 2b and Figure 3(a)(b)(c)).

### 3.5 Effective population size in a haploid-diploid population

We next consider the effective size of haploid-diploid populations with varying degrees of asexuality. Figure 4 plots the effective population size (Eq. 8) relative to the total population size, 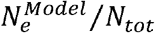, as a function of the frequency of haploids, 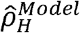 (x-axis), and the class reproductive values (with 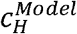 ranging from 0.05 in blue to 0.95 in red). As noted by Bessho and Otto (2017), the effective population size is highest – and drift weakest – at intermediate frequencies of haploids and diploids, which ensures the least sampling error as organisms alternate generations.

**Fig. 4.**
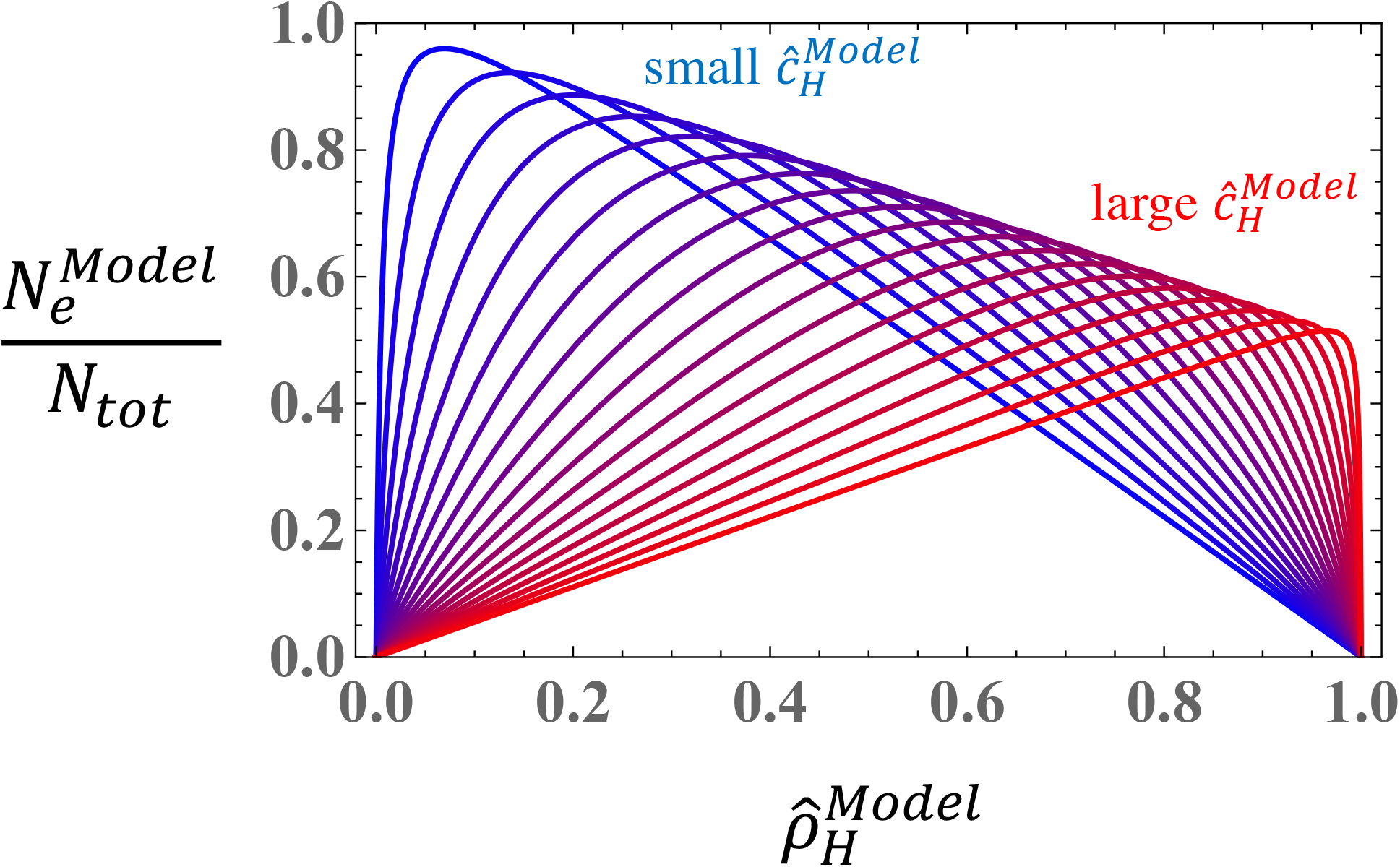
The effective population size of a haploid-diploid population. The relative effective population size over the total population size (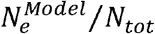, Eq. 7) is shown as a function of the frequency of haploids (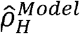, x-axis), when the haploid reproductive value 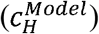 varies from 0.05 (blue) to 0.95 (red) in increments of 0.05. This figure applies to both global and local regulation models.

When haploids and diploids have equal reproductive values, as in the fully sexual case 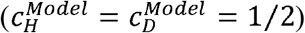, the effective population size is maximized at 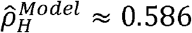. With asexual reproduction, the peak shifts towards whichever ploidy level has the higher reproductive value. For example, if haploids have a high reproductive value (red) then the effective population size is maximized at a higher frequency of haploids, reducing the amount of genetic drift in that phase. Although not illustrated, the peak shifts to 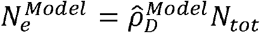 when future populations descend only from diploids 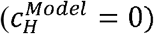 and to 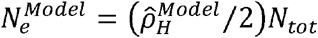 when future populations descend only from haploids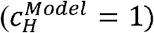, effectively becoming diplontic or haplontic, respectively (with the 1/2 arising because haploids have half the number of chromosomes).

Of course, the reproductive values, as well as the frequency of haploids with global population regulation 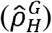, depend in turn on the fertility parameters and the extent of asexuality, as explored in the previous section. Figures 5 (global regulation) and 6 (local regulation) illustrate the effective population size as a function of the frequency of haploid asexuality,*a*_*H*_ (x-axis), and the frequency of diploid asexuality (*a*_*D*_ rising from red to blue), using the parameters in Figures 2 and 3, respectively. The trends are often non-monotonic, with 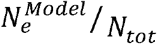 values varying around 1/2 when the parameter values are intermediate. The effective population size is often higher when diploids rarely reproduce asexually (red) rather than when they frequently do (blue), although there are exceptions.

**Fig. 5.**
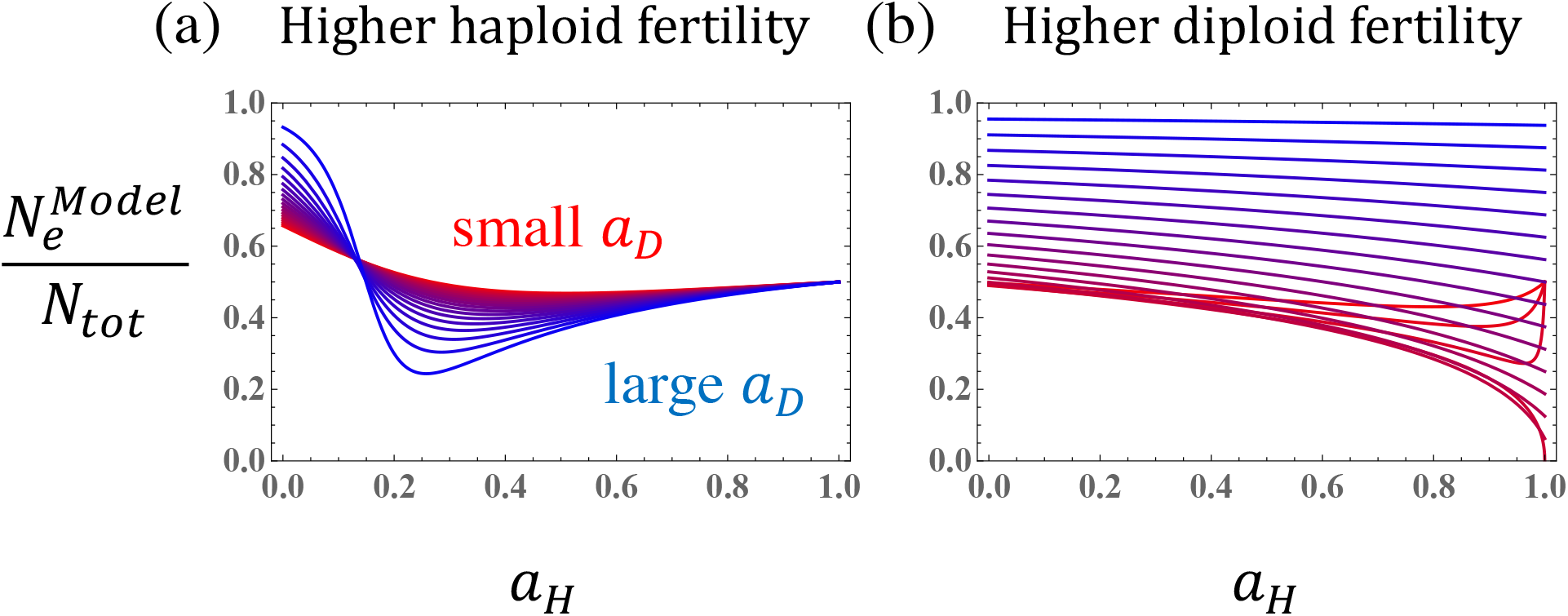
The effective population size of a haploid-diploid population in the global regulation model as a function of the degree of asexuality, *a*_*H*_ and *a*_*D*_. Parameters are the same as in Fig. 2 and determine the relative class reproductive values 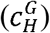 and fraction of haploids 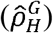 according to Eqs. (2a) and (A.4).

To check whether our analytical result is correct, we simulated a haploid-diploid population for a neutral allele (Figure 7 for the global regulation and Figure 8 for the local regulation). In the classical WF model, the expected heterozygosity within a simulation is 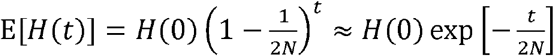. Our simulations reveal that the effective population size predicts the average dynamics of E[*H*_*e*_ (*t*)] (yellow line) where *H*_*e*_ (*t*) 2 *p*_*ave*_ (1 *p*_*ave*_). A slight discrepancy is observed when diploids reproduce primarily asexually (Fig. 7d and 8h), because the assumption that departures from Hardy-Weinberg remain small becomes accurate with drift; nevertheless, our formula for the effective population size continues to perform reasonably well even in this case (especially for the first few hundred generations).

Here, we have tracked the “effective” heterozygosity calculated as *H*_*e*_(*t*) = 2*p*_*ave*_ (1 − *p*_*ave*_) where *p*_*ave*_ is the average allele frequency based on the (typically unknown) class reproductive values. We also find, however, that the effective population size equally well predicts loss of heterozygosity using the naive allele frequency that one might observe by randomly sampling alleles from a population:

*p*_*naive*_ = (2*x*_*MM*_ + *x*_*RM*_ ² *x*_*M*_)/(2*xMM* + 2*x*_*RM*_ + *x*_*RR*_ + *x*_*M*_ + *x*_*R*_) (see Supplementary Mathematica file; Figure 7 and 8). Even the loss of heterozygotes in the diploid population, *x*_*RM*_ / (*x*_*MM*_ + *x*_*RM*_ + *x*_*RR*_ +) is well described by the effective population size derived here.

### 3.6. Fixation probability in a haploid-diploid population

We next compare the above results with numerical simulations estimating the fixation probability of a newly arisen mutation in a haploid-diploid population. When simulating the global regulation model, we assumed that the population has reached the demographic equilibrium, 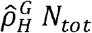 haploids and 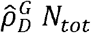 diploids (see Appendix A). We then chose one resident allele *R* at random and replaced it with a mutant allele *M*. After mutation, offspring were sampled from the parental generation according to a multinomial distribution with expected frequencies given by *x*_(*GT*),_ repeating until the mutant allele fixed or was lost from the population. We estimated the fixation probability as the fraction of 100,000 replicate simulations leading to fixation.

We here consider the additive case (*h* = 1/2 and *h*_*e*_ 1/2), where the fixation probability (Eq. 8) simplifies to:

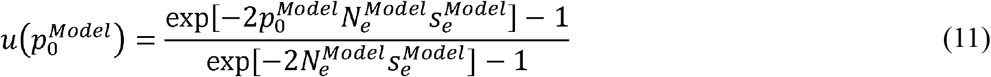

and where 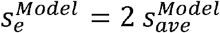 (Eq. 6). Figure 9 plots the fixation probability as a function of the average selection pressure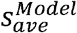, when the reproductive values and chromosome numbers in haploids and diploids are equal (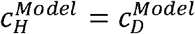 and 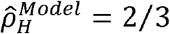) and 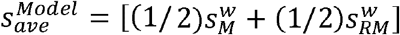. The diffusion Eq. (11) provides an excellent fit, as does the approximation Eq. (9b) for selection coefficients that are positive and not too weak. In this case, the results are the same with global and local population regulation (Fig. 9a and 9b, respectively) and are insensitive to how much selection occurs in the haploid or diploid phases (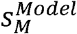 and 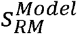, respectively), as long as 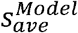 is held constant (see additional simulations in Supplementary *Mathematica* file; Figure 9). As expected, the extent of selection in the haploid versus diploid phase matters more when the mutation is not additive (*h* ≠ 1/2 and *h*_*e*_ ≠ 1/2) (supplementary *Mathematica* file; non additive mutation). Additional simulations are presented in the supplementary *Mathematica* file (Supplementary: Fixation probability for global (or local) regulation model (simulation)).

These confirm that the analytical results remain valid across a broad parameter space (combinations of higher haploid fertility vs diploid fertility, higher haploid asexuality vs diploid asexuality, and higher frequency of haploids vs diploids).

Next, we illustrate the approximate fixation probability, Eq. (9a), as a function of the degree of asexuality (*a*_*H*_ and *a*_*D*_) when the population size is globally (Fig. 10) or locally (Fig. 11) regulated, assuming only selection in haploids or only in diploids. For example, with additive mutations, the fixation probability can be approximated as 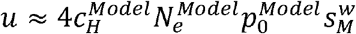 when selection occurs only in the haploid phase or 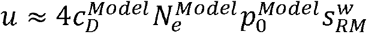 with selection only in the diploid phase, indicating that the fate of mutations depends as much on the strength of selection as on the reproductive value of the ploidy phase in which selection acts (as illustrated in Fig. 2 and 3). Figures 10 (global) and 11 (local) illustrate how the fixation probability depends on the various parameters in the model, particularly the amount of asexual reproduction in haploids (x-axis) and diploids (*a*_*D*_ rising from red to blue). The trends can be understood by the combined effects of the parameters on the reproductive value and the effective population size (e.g., Fig. 11(a) is proportional to the product of Fig. 3(a) and Fig. 6(c)).

**Fig. 6.**
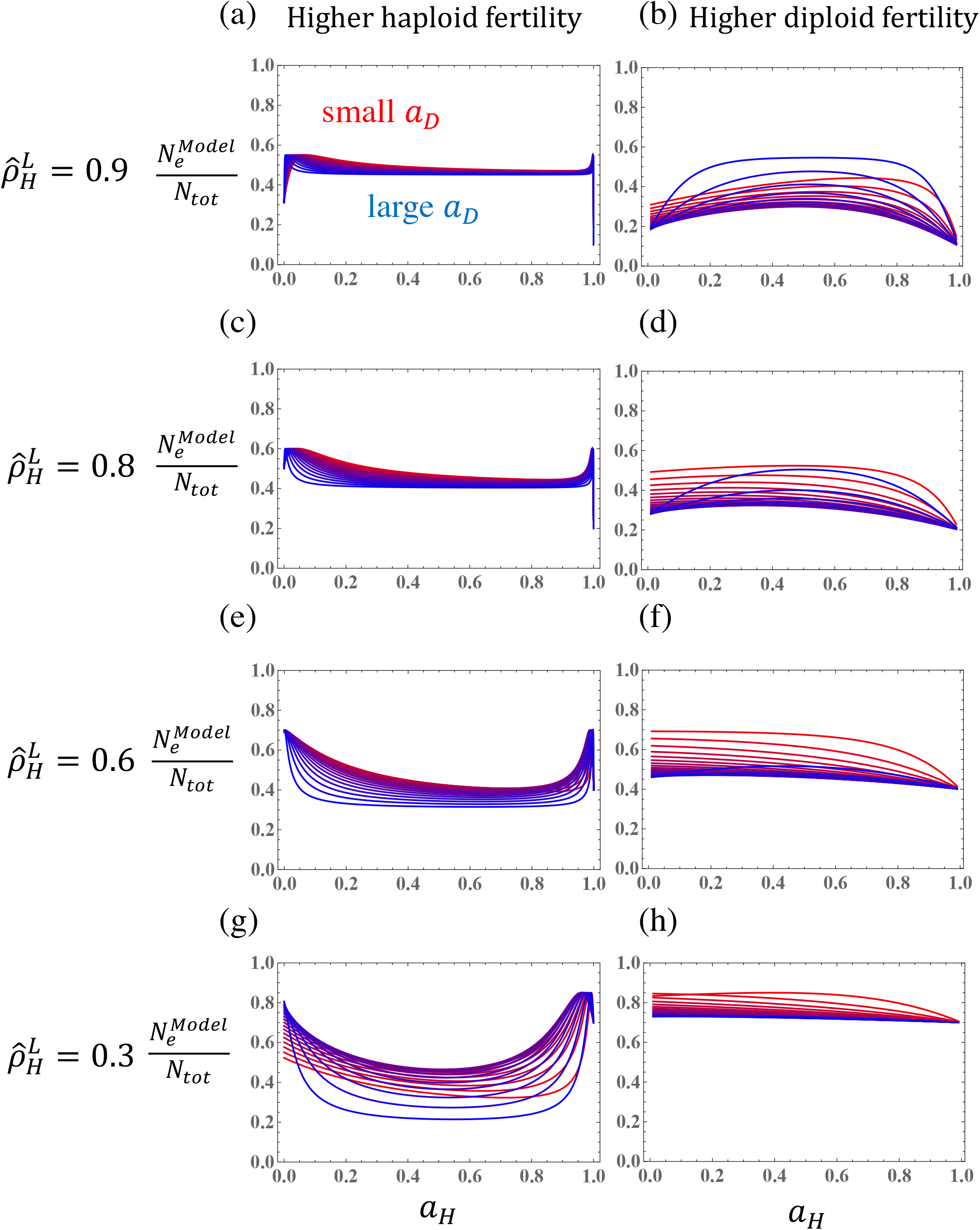
The effective population size of a haploid-diploid population in the local regulation model. Parameters are the same as in Fig. 3 and determine the relative class reproductive values 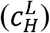 by Eqs. (5). The frequencies of haploids are held fixed at a frequency of (a)(b) 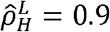, (c)(d) 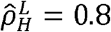, (e)(f) 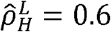, or (g)(h)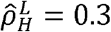.

**Fig. 7.**
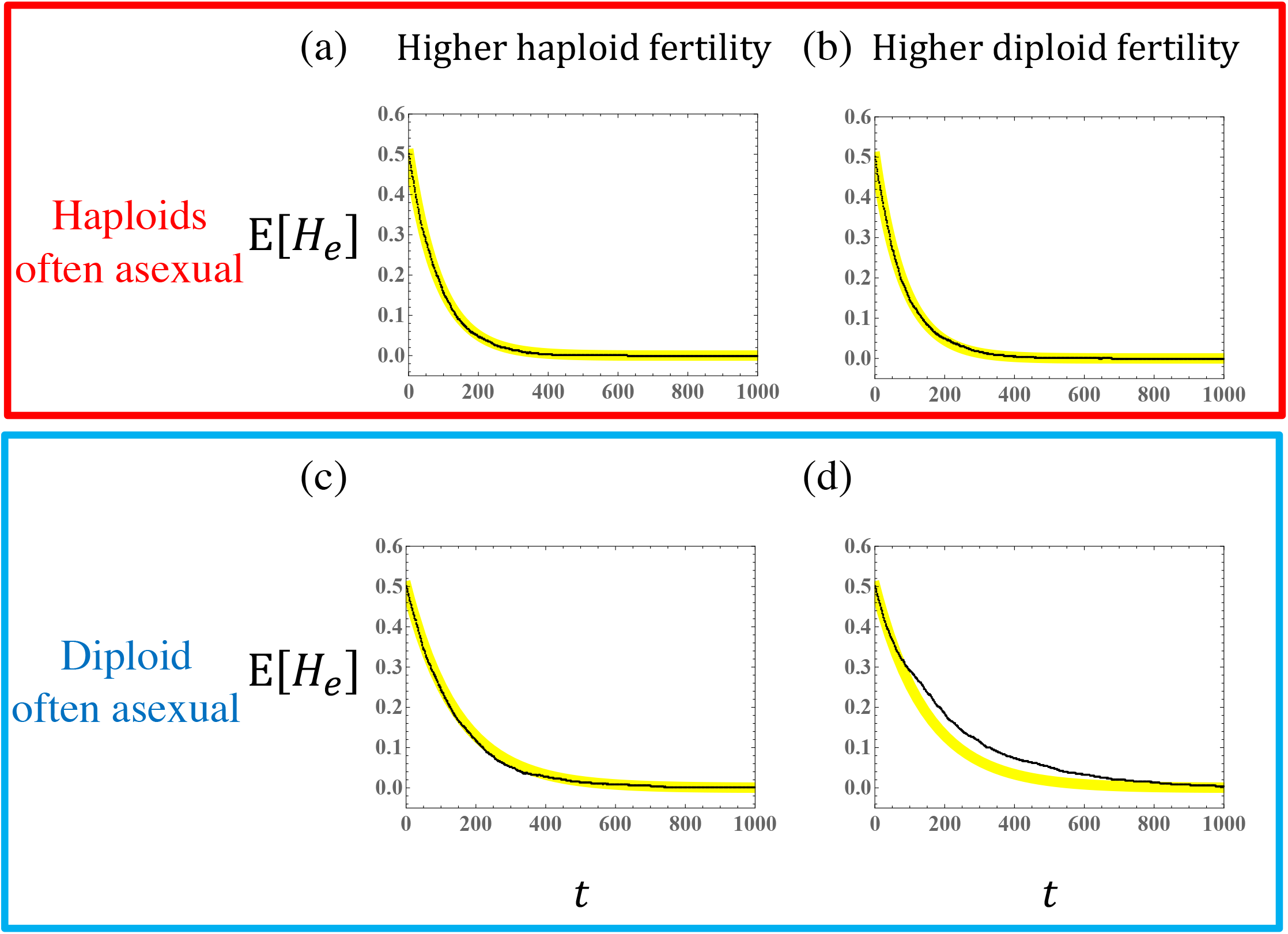
The estimated heterozygosity over time within a globally regulated haploid-diploid population, as measured from 1000 replicate simulations of a neutral model (black dots). The solid yellow curve gives the analytical result for the expected heterozygosity with the effective population size,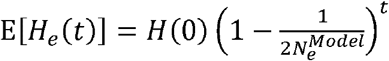. Parameters: *N*_*tot*_ = 90, *f*_*R*_= 0.5. We consider the case when (a)(c) haploids are more fertile than diploids (*w*_*R*_ *=* 5000, *w*_*RR*_ *=* 1000) and when (b)(d) diploids are more fertile than haploids (*w*_*R*_ *=* 1000, *w*_*RR*_ *=* 5000), considering asexual reproduction at a high rate among (a)(b) haploids (*a*_*H*_ *=* 0.8, *a*_*D*_ *=* 0) or (c)(d) diploids (*a*_*H*_ *=* 0, *a*_*D*_ *=* 0.8).

**Fig. 8.**
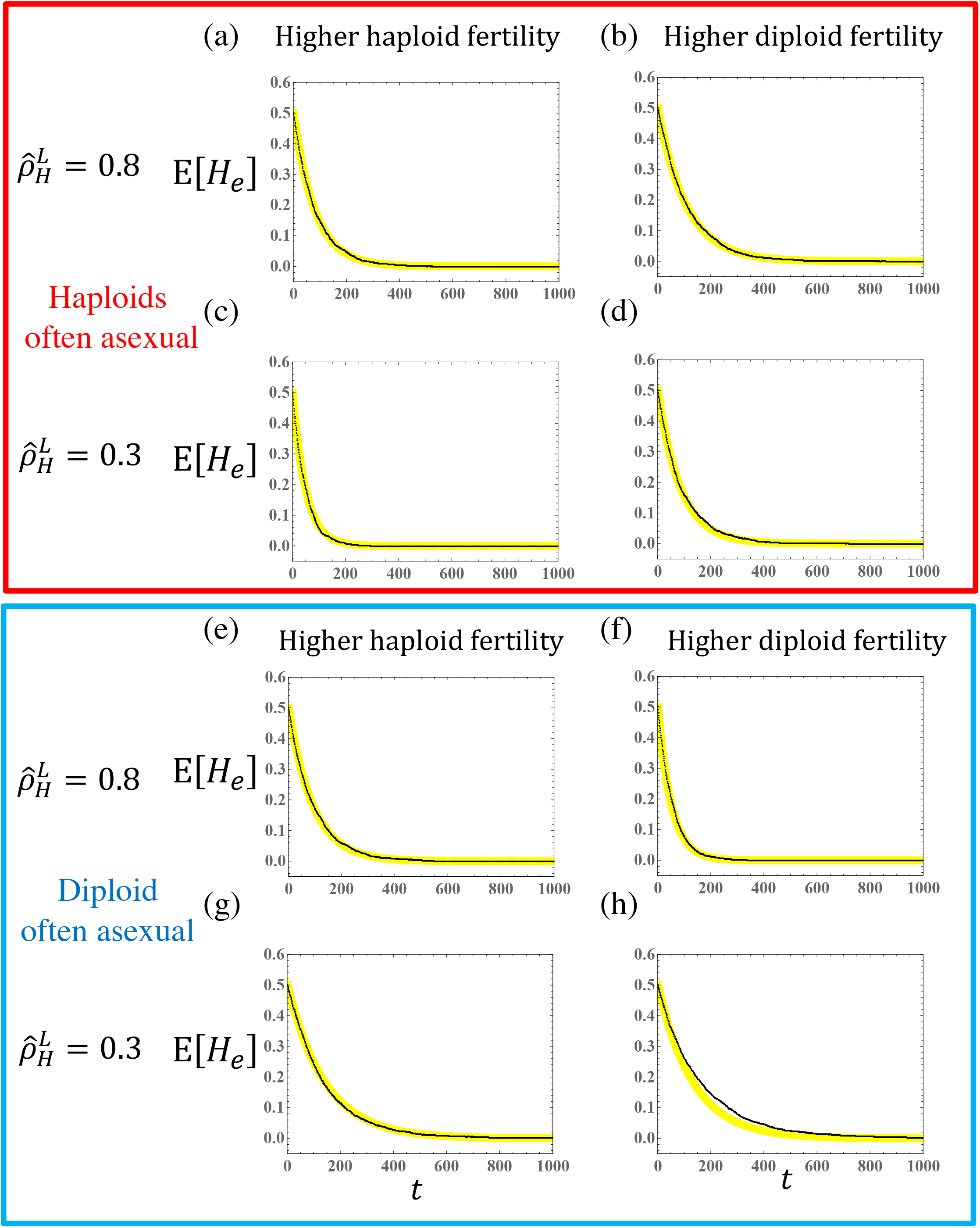
The estimated heterozygosity over time within a locally regulated haploid-diploid population, as measured from 1000 replicate simulations of a neutral model (black dots). The solid yellow curve gives the analytical result for the expected heterozygosity. Parameters are the same as Fig. 7. We consider the case when (a)(c)(e)(g) haploids are more fertile (*w*_*R*_ *=* 5000, *w*_*RR*_ *=* 1000), when (b)(d)(f)(h) diploids are more fertile (*w*_*R*_ *=* 1000, *w*_*RR*_ *=* 5000), when (a)(b)(c)(d) haploids are often asexual (*a*_*H*_ *=* 0.8, *a*_*D*_ *=* 0), when (e)(f)(g)(h) diploids are often asexual (*a*_*H*_ *=* 0, *a*_*D*_ *=* 0.8), holding haploids fixed at frequency of 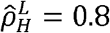 in (a)(b)(e)(f) and at frequency of 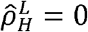 in (c)(d)(g)(h).

**Fig. 9.**
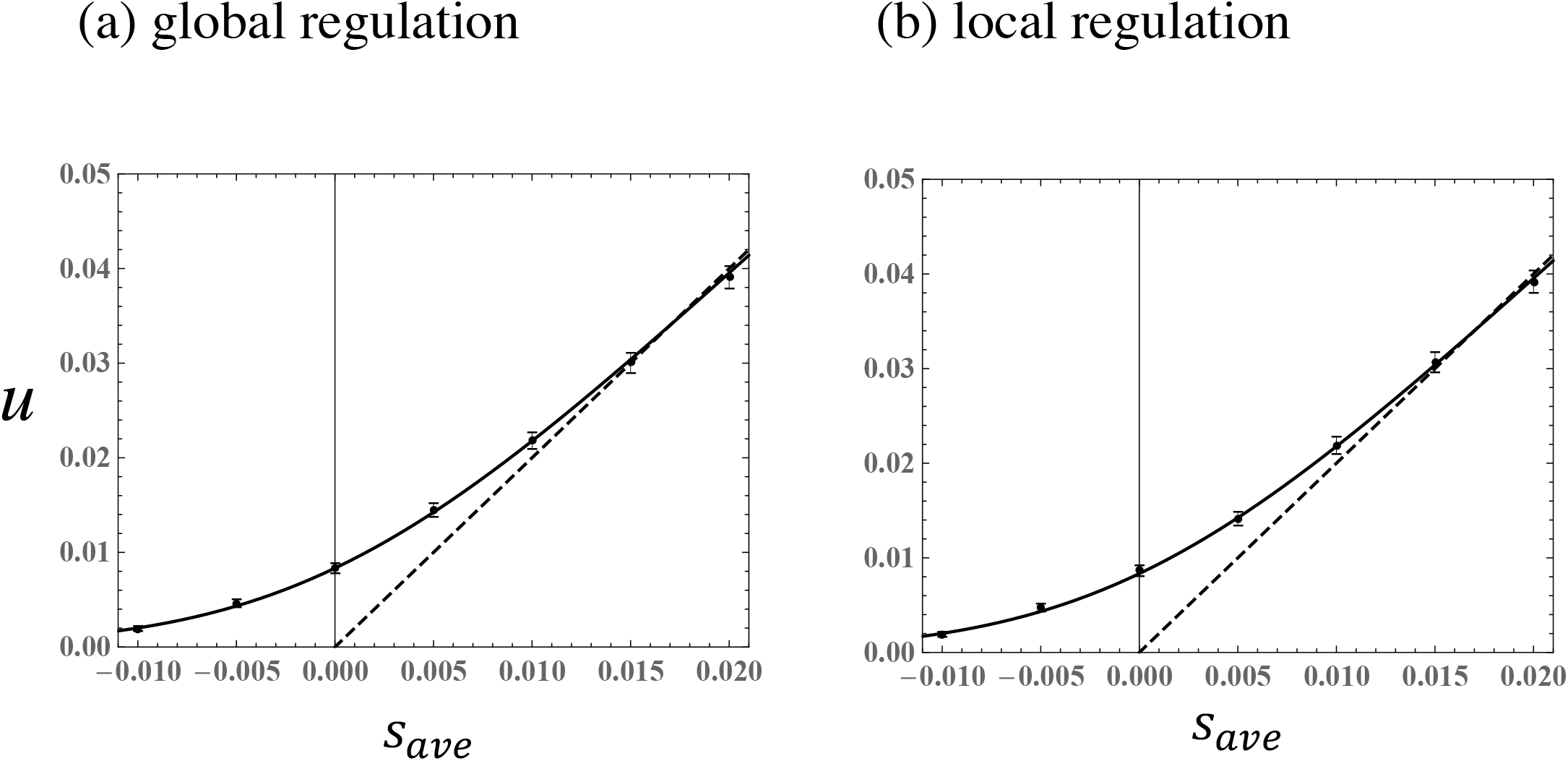
Fixation probability given the average strength of selection, 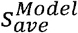, for global (a) and local (b) regulation models. The solid curve gives the analytical result from the diffusion approximation (Eq. (11)) and the dashed curve gives the approximation with weak, additive, and positive selection (Eq. (9b)). Black dots indicate the fixation probability estimated from 100000 numerical simulations with 95% CI (Wilson score interval for binomial). Parameters: *N*_*tot*_*=* 90, *N*_*H*_ *=* 60, *N*_*D*_ *=* 30, *f*_*R*_ *=* 0.5, *w*_*R*_ *= w*_*RR*_ *=* 1000, *a*_*H*_ *= a*_*D*_ *=* 0.1, *h =* 0.5,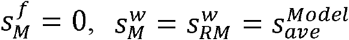, such that the fraction of haploids in the resident population is 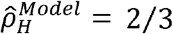 and class reproductive values are equal 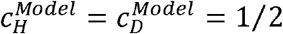. Holding 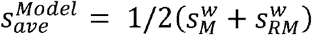 constant, similar results are obtained for a range of different choices of 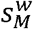and 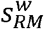 (see supplementary *Mathematica* file).

**Fig. 10.**
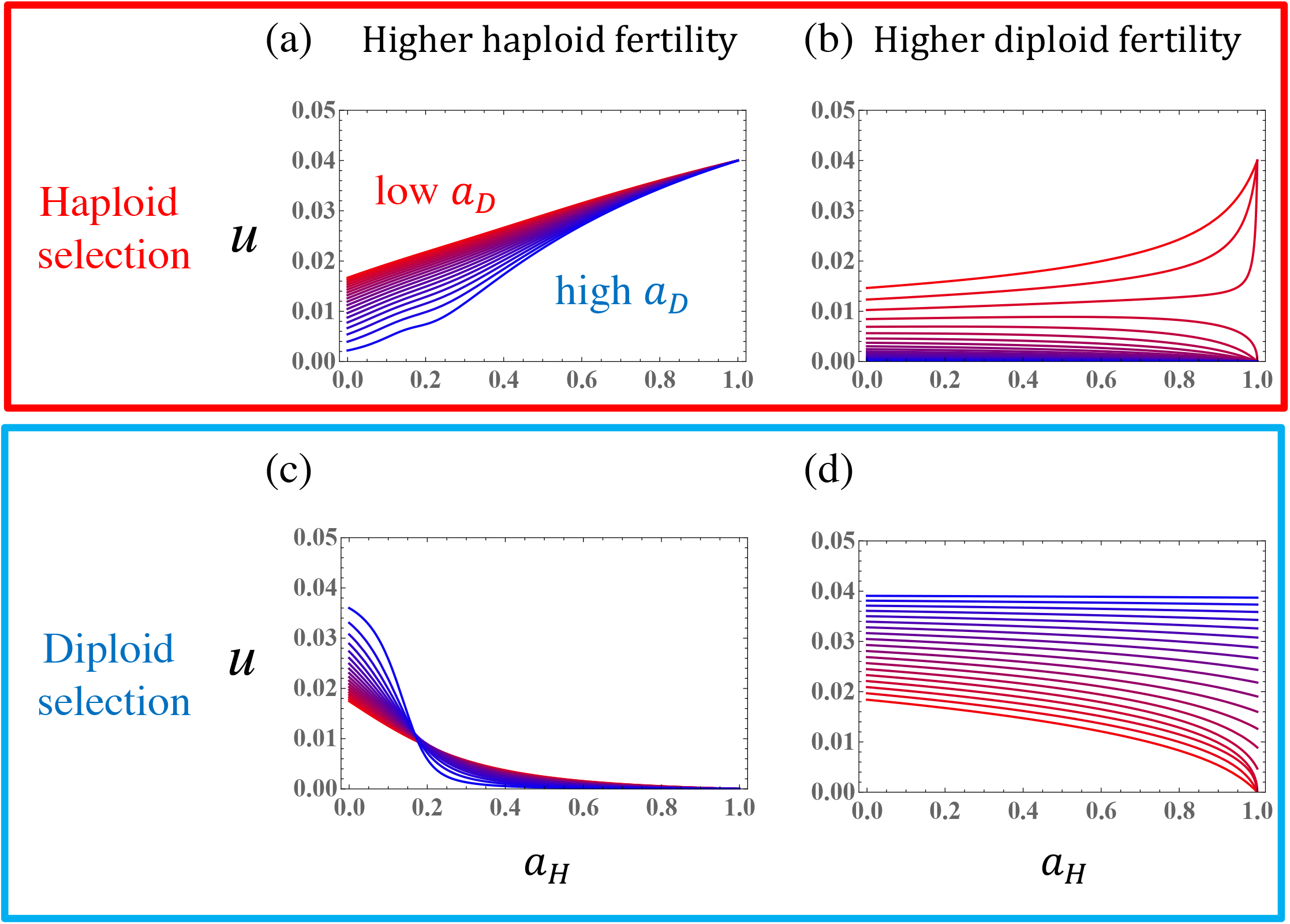
The fixation probability in a haploid-diploid population in the global regulation model. Curves gives the linear approximation for the fixation probability (Eq. (9a)). Parameters are the same as in Fig. 2. Selection acts only in the haploid or diploid phase, with selection coefficients set as 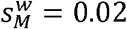 for haploid selection (a)(b) and 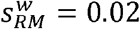 for diploid selection (c)(d).

**Fig. 11.**
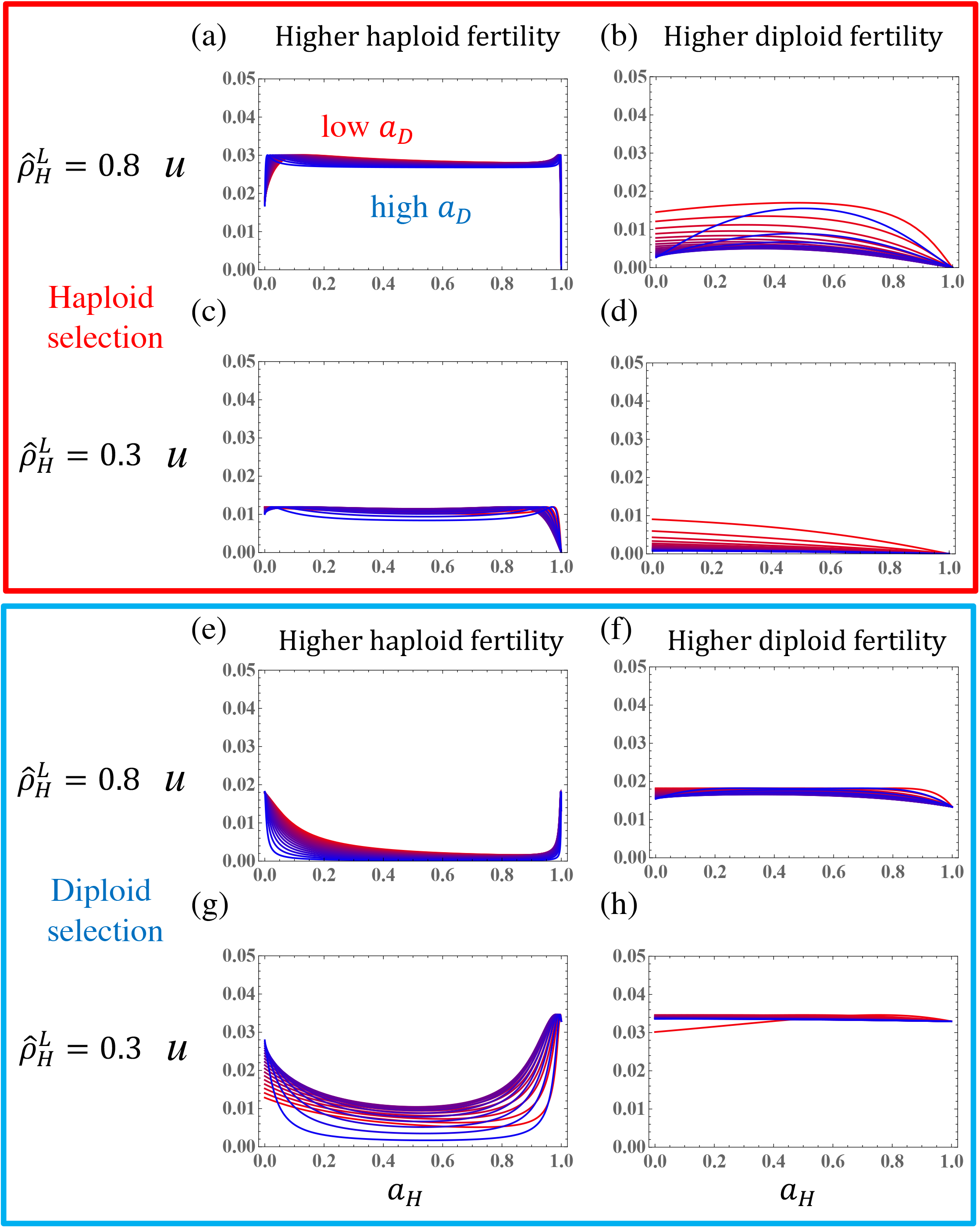
The fixation probability in a haploid-diploid population in the local regulation model. Curves gives the linear approximation for the fixation probability (Eq. (9a)). Parameters are the same as in Fig. 3 and Fig. 10. The frequency of haploids is held fixed at (a)(b)(e)(f) 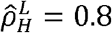, (c)(d)(g)(h) 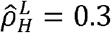. Fertility of haploids is higher than diploids in panels (a)(c)(e)(g), and the opposite condition is considered in panels (b)(d)(f)(h). Selection only occurs in the haploid (a)(b)(c)(d) or diploid stage (e)(f)(g)(h).

## 4. Discussion

Across the phylogenetic tree of life, organisms have diverse and complex reproductive strategies (Bell 1982). Classical population genetic theory has, however, focused most on fully haploid or diploid life cycles with obligate sexuality. In this article we develop a stochastic model for the population genetics of haploid-diploid organisms considering demography, asexuality, and habitat differentiation between haploid and diploid stages. Using a separation of time scales, we derive a diffusion approximation for the change in allele frequency, allowing us to estimate the fixation probability of new mutations, the effective strengths of selection and dominance, as well as the effective population size of haploid-diploid populations.

### 5.1. Natural selection in a haploid-diploid population

Our results indicate that the strength of natural selection and the extent of genetic drift depend strongly on the reproductive value of haploid versus diploid phases. In the simplest case, when the effect of a mutation is weak, additive, positive, and absent in the gamete stage 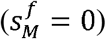, the fixation probability is proportional to the effective strength of selection (Eq. 10), 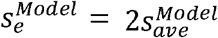, which in turn is proportional to the amount of selection in and the reproductive value of haploids and diploids (Eqs. 2, and 5).

These analytical results reveal some evolutionary principles for populations that undergo an alternation of generations. One consequence is that the balance of opposing selection pressures in haploids and diploids (Eqs. 4 and 6) depends not only on the selection coefficients, but also on the relative reproductive values of haploids 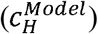 versus diploids 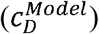. Thus, the very direction of evolution depends on the extent of asexuality in the two phases and the relative survival and fertility of haploids versus diploids when there is “ploidally antagonistic selection” (Immler et al. 2012).

The efficacy of selection to fine tune traits in haploids and diploids also depends on the class reproductive values. For example, when the population is regulated by local density dependence (i.e., the haploid and diploid phases are spatially or temporally distinct), higher reproductive success in haploids increases the efficiency of haploid selection (compare Figure 3a to 3b). However, when there is extremely rare sexuality in haploids (*a*_*H*_ near one), diploid selection tends to be more effective because of increasing competition between offspring from haploids. By contrast, the trends differ with global density dependence (e.g., species that are more isomorphic with small ecological differences between stages). For example, the reproductive value of haploids remains high even when they reproduce primarily asexually in the global regulation model (see Figure 2 when *a*_*H*_ approaches one), because haploids then make up a larger proportion of the total population size (see inset figures). Thus, whether selection is effective in the haploid phase when that phase mainly reproduces asexually is quite sensitive to the nature of competition.

Our work can also be useful in the design of field studies and the interpretation of data for species that alternate generations. To understand the efficiency of selection on haploid and diploid phases, we not only need data about the fraction of haploids and diploids and their fertility and mortality (e.g., Thornber and Gains 2004; Vieira et al., 2018a; Vieira et al. 2018b), but we also need to know about the extent of asexuality in each phase and whether they compete for common or different resources.

### 5.2. Genetic drift and effective size

The impact of random genetic drift on the genetic diversity of haploid-diploid population depends on the effective population size (Eq. 7). As we had found previously in a haploid-diploid model with obligate sexuality (Bessho and Otto 2017, pp. 431), the effective population size with asexuality is generally smaller than the total number of individuals and again depends strongly on the reproductive value of each phase (Figures 4-6). With obligate sexuality, the reproductive values of haploids and diploids are equal, drift is minimized for a new allele (the effective population size times *p*_0_ is maximized) when haploids comprise 2/3 of the population, making the number of chromosomes equal between haploids and diploids. Asexual reproduction, however, causes the reproductive value of haploids and diploids to differ (Eqs. 2 and 5). Consequently, drift is lessened if the phase with the higher reproductive value is more common (see shifts in peaks in Figure 4).

### 5.3. Evolution in haploid-diploid populations with asexuality

We find that evolution in haploid-diploid populations is substantially different than either fully haploid or fully diploid populations and also substantially different than fully sexual haploid-diploid populations. The model considered in this paper allows selection in both haploid and diploids phases but does not require a strict alternation of generations, allowing individuals to produce offspring of either the same ploidy type (asexually) or the opposite ploidy type (sexually).

Here we consider the implications for evolution, focusing on the simplest case of additive selection with weak selection, where the fixation probability is given by Eq. (9a). This formula indicates that evolution in a haploid-diploid population is dominated by (i) the average selection across haploid and diploid stages, (ii) the ploidy ratio in the population, and (iii) the class reproductive values of haploids and diploids. Comparing this probability with the classical result 2*s* reveals that evolution in a haploid-diploid population differs substantially in three ways from evolution in a fully haploid (or diploid) population.

First, and most obvious, evolutionary changes are shaped by selection in both haploid and diploid stages. As we see in subsection 5.1, because the reproductive values of haploids and diploids determine how selection in each phase contributes to the effective selection coefficient (6a), selection is not a simple average across ploidy stages but depends on the details of how the species reproduces. Second, the ploidy ratio of haploids versus diploids strongly affects evolution. In particular, because genetic drift is strongest when either haploids or diploids are rare (Figure 4), the effective size dramatically decreases when the ploidy ratio is strongly skewed (i.e., most individuals are haploids or diploids; Figure 4), reducing the efficacy of selection. Third, the reproductive values of haploid and diploid stages depend in a complex way on selection in the two phases and their mode of reproduction (Figures 2-4), affecting the course of evolution through both selection and genetic drift.

Evolutionary dynamics in completely sexual haploid-diploid populations is substantially simpler. In particular, with a strict alternation of generations (no asexuality), the class reproductive values of haploids and diploids are equal (Bessho and Otto 2017). As a consequence, selection across the life cycle, is half the strength of selection in haploids and half in diploids, regardless of the frequency of the two types within the population.

With both asexual and sexual reproduction, however, the average selection, the ploidy ratio, and the reproductive values of haploids and diploids all depend on the fertility parameters and the degree of asexuality. The resulting interplay alters the fixation probability of new mutations, as illustrated in Figures 10 and 11. While the interplay is complex and often non-monotonic, some patterns emerge.

An important take-home result is that the form of competition acting to regulate population size strongly affects the reproductive values of haploids and diploids (Eqs. (2) and (5)) and thus dramatically alters the chance that a new beneficial mutation fixes (equation 8; Figures 10 and 11). We contrast two forms of population regulation: global, where haploids and diploids are competitively equivalent, and local, where only individuals of a given ploidy level compete with one another.

With global regulation, the fixation probability for an allele that benefits the haploid phase is generally higher when haploids are more asexual and diploids more sexual (Figure 10), increasing the fraction of haploids in the population (Figure 2), but there are exceptions when diploids have higher fertility and reproduce predominantly asexually (bluer curves in Figure 10b). Similarly, the fixation probability for an allele that benefits the diploid phase is generally higher when diploids are more asexual and haploids more sexual (Figure 10), increasing the fraction of diploids experiencing selection, with exceptions when haploids have higher fertility and are predominantly asexual (right hand side of Figure 10c). In summary, selection in a particular ploidy phase is typically more effective when that phase is more common and tends to breed true (asexual reproduction), with exceptions when the other ploidy phase is a greater evolutionary “source” of future generations, with a higher reproductive value (higher fertility and more asexual). These exceptions occur because passage through the other ploidy phase then becomes essential for an allele to avoid loss while rare, even when selection does not act in the other phase.

The patterns with local regulation of population size differ substantially. Because local population regulation by ploidy level determines the equilibrium fraction of haploids and diploids, the fixation probability tends to be less dependent on the exact degree of asexuality, as long as there is some sexual and some asexual reproduction (flatter curves with more overlapping red/blue in Figure 11). In this case, the fixation probability of an allele favoured in the haploid phase is higher if haploids are more fertile (Figure 11 panels a and c versus b and d) and if haploids are more common (panels a and b versus c and d), with the reverse holding for alleles favoured in the diploid phase. When populations are predominantly sexual or predominantly asexual, however, there can be rapid changes in the fixation probability (Figure 11), resulting from rapid changes in the reproductive value of each phase (Figure 3).

### 5.4. “Ploidally-structured” populations

The key role that reproductive values play in this work is analogous to the role that patch dynamics play in two-patch models of evolution. In a spatially structured population, subdivided local populations are genetically connected by migration. A haploid-diploid system can be seen as being ploidally structured, where gene flow describes the movement of alleles through sexual reproduction, with meiosis causing flow to haploidy and syngamy flow to diploidy. We note that our research reveals that all qualitative results are equally accurate for evolution in a two-patch system (see Supplementary *Mathematica* File; Comparison with two-patch system). For example, fixation probability strongly depends on class reproductive values of each patch.

This analogy suggests an interesting idea: complex reproductive systems can be considered and analyzed using the tools of metapopulation theory. For example, many eukaryotes including terrestrial plants, insects, and fishes, often exhibit ploidy variation, including polyploid members (Otto and Whitton 2000; Comai 2005). In such species, individuals characterized by different numbers of chromosomes coexist, with complex reproductive relationships causing gene flow between them (Ramsey and Schemske 1998). Similarly, social insects often exhibit complex sex determination systems linked with ploidy levels (haplodiploidy).

Our research suggests that these ploidally-structured populations can be fruitfully treated as metapopulations. Selection and drift in populations with diploids, triploids, and tetraploids can, for example, be considered as a three-patch model. In this system, we conjecture that the average strength of selection that is evolutionarily relevant would be the mean selection coefficient in each ploidy class, weighted by its class reproductive value, with additional terms coming from reproductive interactions (akin to the term of 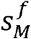 in Eqs. 4).

Many evolutionary aspects of haploid-diploid populations remain to be investigated. One avenue that we are exploring is how model parameters can be estimated from field data. For example, the analogy between spatially and ploidally structured population suggests that genetic differences between haploids and diploid can be used to estimate gene flow between them (i.e., rates of sex), akin to using Fst to inform estimates of migration (e.g., Slatkin 1987). Another fruitful avenue for further work is to determine how fluctuations in population size affect the effective population size of species that alternate generations. In classical population genetics theory, such fluctuations can be captured by using the harmonic mean population in place of the total population size (Karlin 1968). It is unclear, however, whether the same is true in haploid-diploid populations. Can the harmonic total population size simply replace *N*_*tot*_ in the global model of population regulation? Similarly, can the harmonic population sizes of haploids and diploids replace *N*_*H*_ and *N*_*D*_ with local regulation? The answer is unclear because population size fluctuations perturb the fast ecological dynamics away from the steady state (especially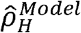), and the impact of these perturbations on selection and drift is unknown. Further research is needed to clarify evolutionary processes in the wide variety of species that alternate generations.

## Supporting information

Supplementary_Mathematica_file

## Acknowledgements

We here thank Alireza Ghaseminejad for help in developing our manuscript. We also thank H. Ohtsuki and Y. Uchiumi for the helpful comment.

## Appendix A. Fixation probability in a haploid-diploid Wright-Fisher model using a diffusion approximation

### A.1. Equilibrium with global regulation

We derive the fixation probability in a haploid-diploid population using a diffusion approximation (e.g., Bessho and Otto 2017). We first derive the stable equilibrium in the global regulation model, allowing for asexual reproduction in each phase. In the Wright-Fisher model, all individuals reproduce and then the parents die (non-overlapping generations). Let *b*_(*GT*)_ represent the number of reproductive cells of each type in the next generation:

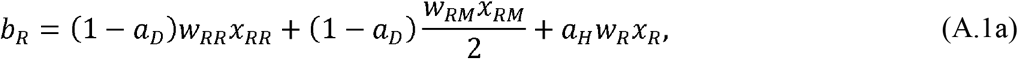

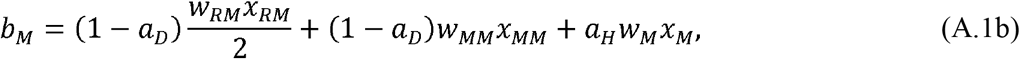

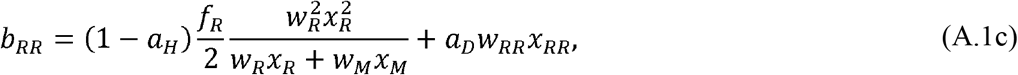

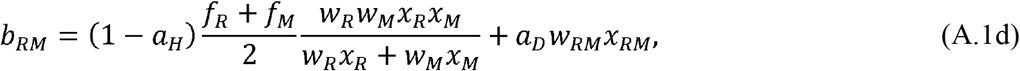

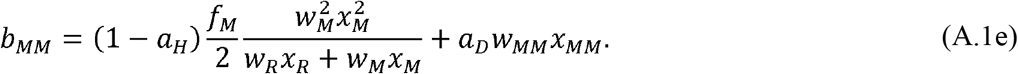

The probability that a reproductive cell of genotype (*GT*) is sampled from the offspring produced by the previous generation of adults is

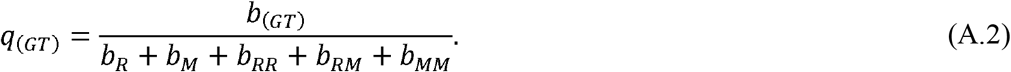

Therefore, the composition of offspring in the next generation is given by the multinomial distribution, sampling *N*_*tot*_ individuals in proportion to Eq. (A2). Using Eq. (A1) and (A2), we describe the conditional expectation of change in the number of individuals of genotype (*GT*), Δ *x*(_*GT*_) (*t*) = *x* (_*GT*_) (*t* + 1)− *X* (_*GT*_)(*t*), as

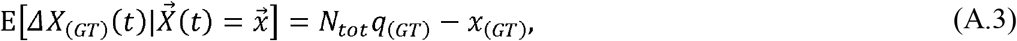

Where 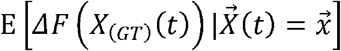 is the conditional expected value for change in the function *F* of the random variable given that 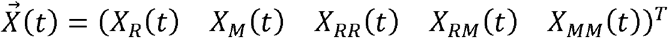 equals 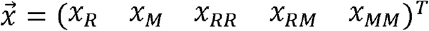.

To simplify this fully stochastic system, we assume that the resident population is large and treat demographic changes deterministically prior to the appearance of the mutation. Considering the dynamics of the resident population, we then find the equilibrium of these dynamical equations by solving 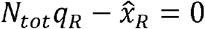 and 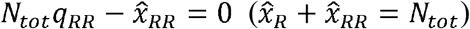. Setting 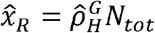 and 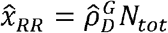, the fraction of haploids 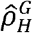 (and diploids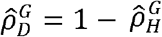) at equilibrium becomes,

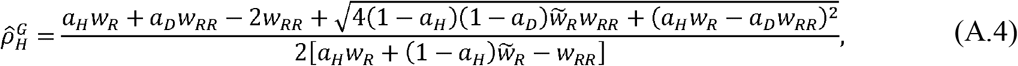

where 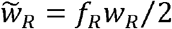 is the fertility of haploids considering the cost of sex (see Supplementary *Mathematica* file for the step-by-step derivation; Supplementary parts). We note that, when the fertility of haploids is much greater than that of diploids (*w*_*R* ≫_*w*_*RR*_), the frequency of haploids in a population approaches *a*_*H*_ /{*a*_*H*_ + [(1 − *a*_*H*_) *f*_*R*_ /2]}.which is less than one because sexual reproduction of the haploids produces diploids (the (1 − *a*_*H*_) *f*_*R*_ /2term). Conversely, when the fertility of diploids is much greater than haploids(*w*_*R ≫*_*w*_*RR*_), approaches 1 − *a*_*D*_, the rate at which diploids undergo meiosis.

### A.2 First moment of change in allele frequency

To derive the first moment of change in allele frequency, *m* ^*Model*^ *(p*_*ave*)_, we apply a separation of time scales (e.g., Nagylaki 1976; Otto and Day 2007; Bessho and Otto 2017). Details of the calculation are represented in the Supplementary *Mathematica* file (Diffusion approximation for the global (local) regulation model). We first transform the expected change in the number of individuals of each type (five variables that sum to*N*_*tot*_) into the expected change in a new set of four variables, *Θ* ∈ {*p*_*ave*_, *δ*_*p*_, *η*_*HW*_, *ρ*_*H*_}, described by the functions:

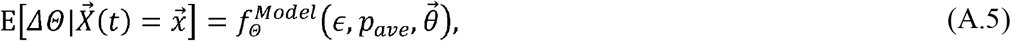

where 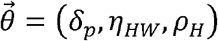 and *∈* is proportional to the selection coefficients and assumed small (the functions *f* are given explicitly in the Supplementary *Mathematica* file). With local regulation, *ρ*_*H*_ is assumed fixed at *N*_*H*_/*N*_*tot*_ and dropped from the variable set,. *Θ*.

To constant order (setting the small changes due to selection to zero, *ϵ* →0), the fast ecological dynamics of the system are described by: 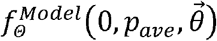. This system of equations rapidly approaches a steady state found by solving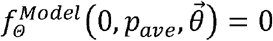, which gives 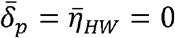, and 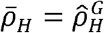 (Eq. A.4). To this order, the steady state change in allele frequency is zero,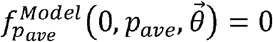. We then describe slower changes, including changes in allele frequency due to selection, by describing the deviations that occur around this steady state. Specifically, to order *∈*, the variables are allowed to deviate from the steady state by 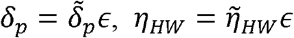, and 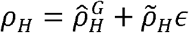 and the dynamics 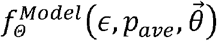, and are then approximated using a Taylor series expansion. Defining the average allele frequency by combining haploid and diploid populations using an arbitrary weighting, *p*_*ave*_ = *ωp*_*H*_(1 − *ω*)*p*_*D*_, we show in the Supplementary *Mathematica* file (Class reproductive value in Supplementary parts) that setting the weights proportional to the class reproductive values (given by Eq. (2) with global regulation and Eq. (5) with local regulation) is the only choice that separates evolutionary change in *p*_*ave*_ from changes in the other variables to order. ϵDefining the average allele frequency in this way (Eq. 1a), taking the Taylor series, and reporting the results in the initial parameters (e.g.,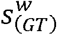 rather than 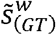, where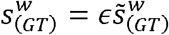) the change in allele frequency becomes:

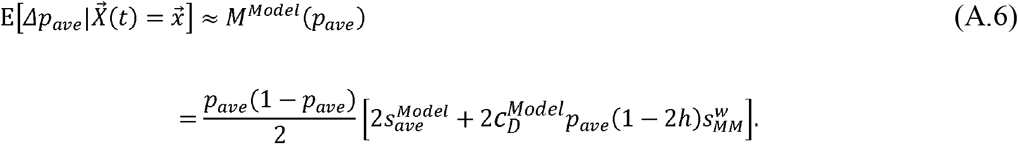

In the supplementary Mathematica file, we represent an example of the simulation of 4-dimensional dynamics calculating the fraction of times that the mutation fixed in the population (see section “Example of 4-dimensional dynamics”)

### A.3 Second moment of change in average allele frequency

We next derive the second moment of change in average allele frequency in a haploid-diploid population with asexuality. We again assume that the population size is very large, that selection is very weak, and that the system has approached the steady state in (*δ*_*p*_, *η*_*HW, ρ H*_), ignoring deviations that are of 0 (ϵ) Because selection is assumed weak, the second moment is well approximated by that of the neutral model (to constant order, ϵ →0).

Under these assumptions, the fraction of haploids in a population is relatively fixed in both the global and local regulation models, and we can sample the haploid offspring according to a binomial distribution, with expectation and variance: 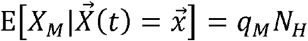 and 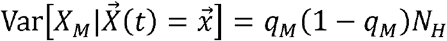 Where *q*_*M*_ = *b*_*M*_ /(*b*_*R*_ + *b*_*M*_) To simplify the equation, we set 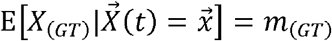 and 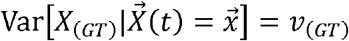, finding that:

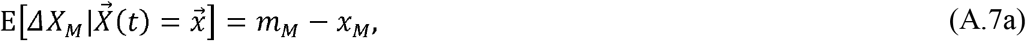

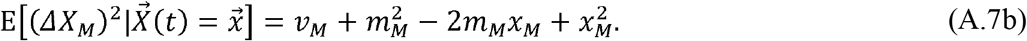

In terms of allele frequencies (rather than numbers), we have the first and second moments for the haploid offspring population, 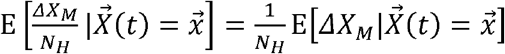 and 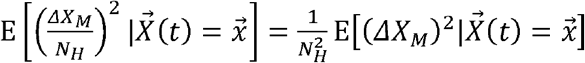.

Similarly, the diploid offspring are sampled according to a trinomial distribution, with expectation, variance, and covariance: *m*(_*GT*)_ =*q*_(*GT*)_*N*_*D*,_ *v*_(*GT*)_= *q*_(*GT*)_ (1 − *q*_(*GT*)_*N*_*D*_, and 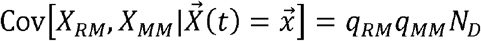, where *q*_(*GT*)_= *b*_(*GT*)_/(*b*_*RR*_ + *b*_*RM*_ + *b*_*MM*_) To derive the moments of the allele frequency in diploids, we define, *y*_*M*_=(*x*_*RM*_/2) + *x*_*MM*_ and *Y*_*M*_= (*X*_*RM*_/2) + *X*_*MM*_. The moments of random variable *Y* are then:

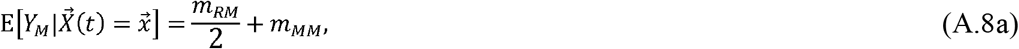

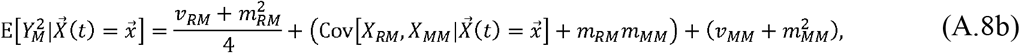

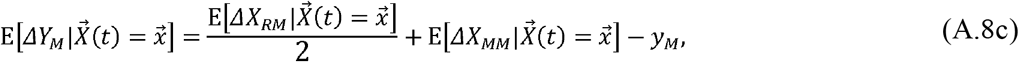

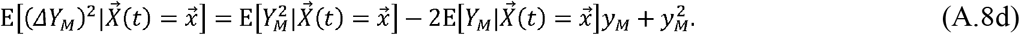

To consider the change in average allele frequency across the entire population, we difine 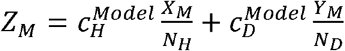 and consider the expectation of change in this random variable. Plugging in Eqs. (7a), (7b), (8c), and (8d), we have

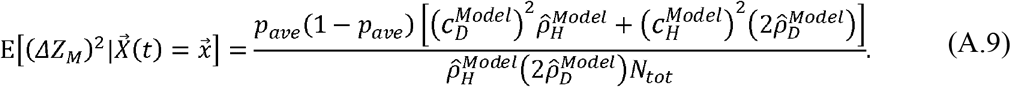

After transforming time scales using the variable τ = *t/ N*_*tot*_ and defining, *P* (*τ*) =Z (*N*_*tot*_τ), we have the diffusion coefficient 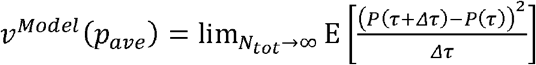 by taking the limit *N*_*tot →*_, ∞ giving Eq. (3c). Similarly, we derive the drift coefficient using Eq. (A6) *m* ^*Model*^ = M^*Model*^ *N*_*tot*_), giving Eq. (3b).

### A.4 Class reproductive value of global regulation model

We now calculate the reproductive value of being haploid or diploid, assuming only resident individuals. Specifically, we are interested in knowing what the long-term contribution of an individual is to the future population if it is sampled from either haploids or diploids. Here, we describe the population dynamics in a haploid-diploid population of resident alleles,

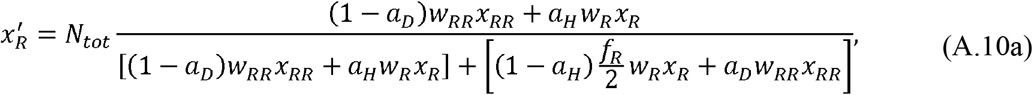

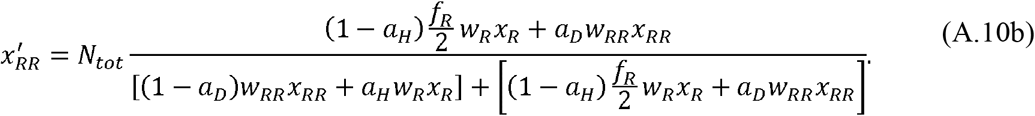

where, 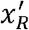 and 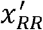 indicate the average number of haploids and diploid at the next time step.

The reproductive value of an individual type is generally calculated for linear models without density dependence. Because the dynamics of *x*_*R*_ and *x*_*RR*_ converge to equilibrium, we modify the recursions so that the total offspring pool size (denominator) is held constant at its steady state value. This gives us a linear model that allows us to focus on the effects of individual fitness and movement without the density-dependent feedback:

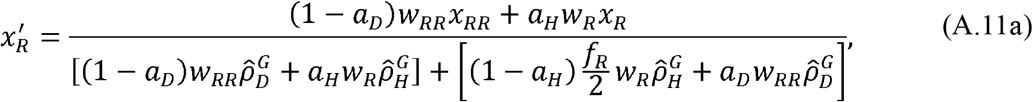

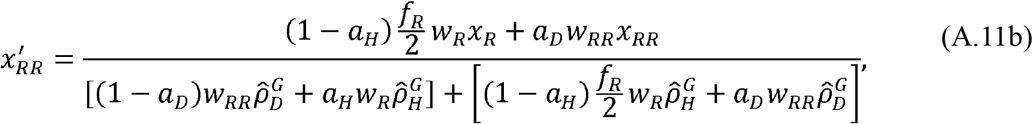

Eqs. (A.11) can be described by the matrix, 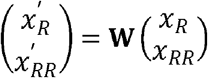. At demographic equilibrium, the leading eigenvalue of matrix **w** should be one (see details in Supplementary*Mathematica* file; Supplementary parts in “Diffusion approximation for the global regulation model”). The left eigenvector associated with this eigenvalue is:

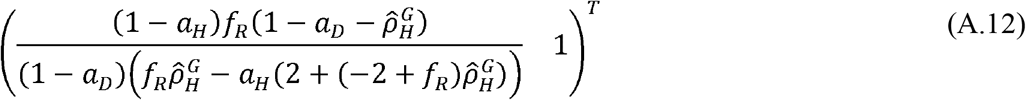

which gives the relative individual reproductive value for a single individual haploid or diploid. The class reproductive values scale this up the contribution of each class,

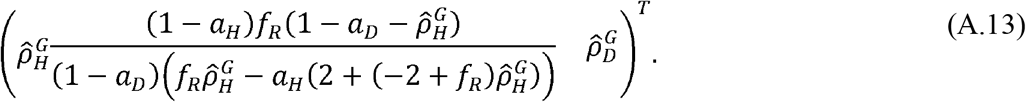

These are proportional to the class reproductive values, Eqs. (2).

### A.4. Class reproductive value of local regulation model

We next calculate the reproductive value of being haploid or diploid with local regulation. We describe the population dynamics in a haploid-diploid population of resident alleles,

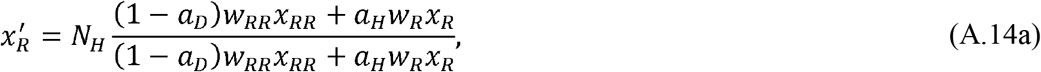

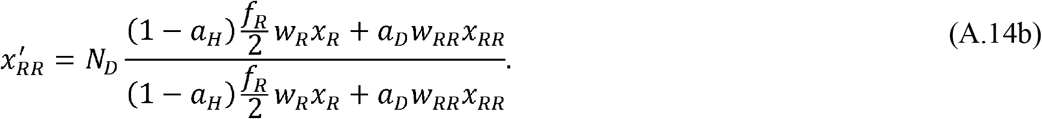

For the local regulation model, because both the haploid and diploid population sizes are fixed, we have the trivial dynamics 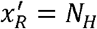 and 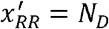.

To assess the impact of slight changes around this state, we again hold the size of the offspring pool constant at its steady-state value, obtaining the linear set of equations:

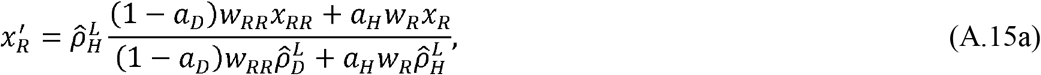

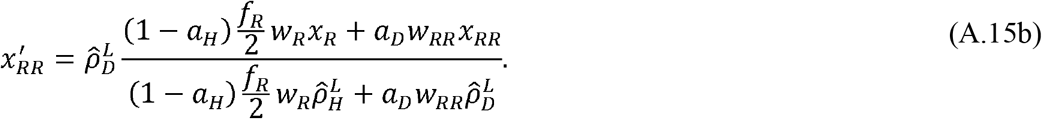

Eqs. (A.15) can be described by matrix, 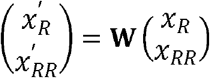. The leading eigenvalue of **W** is again one because of the assumption that the population is at its steady-state size (see details in Supplementary *Mathematica* file; Supplementary parts in “Diffusion approximation for the local regulation model”), and the associated left eigenvector becomes,

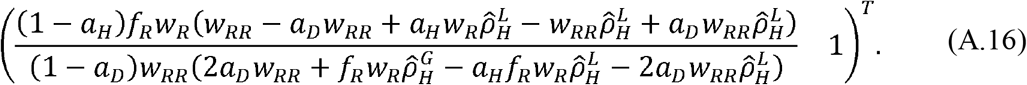

This vector gives the relative reproductive value for a single individual that is either haploid or diploid. The class reproductive values scale this up, with the contribution of each class being proportional to the class reproductive values, Eqs. (5).

## Supporting information

S1. Supplementary *Mathematica* file.

